# Quantitative Structural Assessment of Graded Receptor Agonism

**DOI:** 10.1101/617100

**Authors:** Jinsai Shang, Richard Brust, Patrick R. Griffin, Theodore M. Kamenecka, Douglas J. Kojetin

## Abstract

Ligand-receptor interactions, which are ubiquitous in physiology, are described by theoretical models of receptor pharmacology. Structural evidence for graded-efficacy receptor conformations predicted by receptor theory has been limited, but is critical to fully validate theoretical models. We applied quantitative structure-function approaches to characterize the effects of structurally similar and structurally diverse agonists on the conformational ensemble of nuclear receptor peroxisome proliferator-activated receptor gamma (PPARγ). For all ligands, agonist efficacy is correlated to a shift in the conformational ensemble equilibrium from a ground state towards an active state, which is detected by NMR spectroscopy but not observed in crystal structures. For the structurally similar ligands, ligand potency is also correlated to efficacy and conformation, indicating ligand residence times among related analogs can influence receptor conformation and function. Our results derived from quantitative graded activity-conformation correlations provide new experimental evidence and a platform with which to extend and test theoretical models of receptor pharmacology to more accurately describe and predict ligand-dependent receptor activity.

## INTRODUCTION

Receptor theory has been used to describe the actions of pharmacological ligands in various forms for nearly a century (Maehle et al. 2002, Kenakin 2004). The idea of a receptor has evolved from a conceptual “black box” to one founded in the principles of biophysics and allostery (Changeux 2012). The two-state model of receptor activation (Leff 1995), which extended the Black/Leff operational model of pharmacological agonism (Black et al. 1983) with the Monod-Wyman-Changeux (MWC) model of protein allostery (Monod et al. 1965) to describe the actions of pharmacological receptor ligands, conceptually represents a minimal theoretical model to describe the action of ligands within the context of a binary ligand-receptor complex. More complex models were subsequently developed that accounted for improved understanding of receptor functions. The extended ternary complex (ETC) model describes how the receptor-ligand complex influences interaction with an effector protein or a signaling pathway (Samama et al. 1993), whereas the cubic ternary complex (CTC) model extends the ETC model to account for receptor-effector interactions in the absence of ligand (Weiss et al. 1996). These and other theoretical receptor models could be further improved with a more comprehensive experimental understanding of how ligands affect receptor structure and function.

Applying theoretical receptor models to graded activity dose-responsive pharmacological data is common in studies of membrane receptors, including G protein-coupled receptors (GPCRs), ligand-gated ion channels, and enzyme-linked receptors (Kenakin 2004). These receptors bind extracellular ligands and transduce signals across the cell membrane via conformational rearrangement of the intracellular portion of the membrane receptor to affect various downstream signaling pathways, the activities of which can be measured in cellular assays and applied to pharmacological models of receptor function. Conceptually, the principles of receptor theory also apply to nuclear receptors, a superfamily of intracellular transcription factors that recruit chromatin remodeling transcriptional machinery in a ligand-dependent manner to control gene expression (Weikum et al. 2018). Nuclear receptor agonists, which bind to an internal hydrophobic orthosteric pocket within nuclear receptor ligand-binding domain (LBD), activate transcription by stabilizing structural elements that comprise the activation function-2 (AF-2) coregulator interaction surface, located adjacent to the ligand-binding pocket, including helix 3, helix 4/5, and a critical regulatory switch element, helix 12. Agonists stabilize an active AF-2 surface conformation, which increases the binding affinity for and recruitment of transcriptional coactivator proteins that in turn promotes chromatin remodeling and increased transcription. Although receptor theory, as practiced in the membrane receptor fields, is not used in the nuclear receptor field, in principle the functional endpoints derived from nuclear receptor functional assays can be applied to the same pharmacological models of receptor function.

It has been challenging to structurally observe the graded receptor activity conformations predicted by receptor theory. Although ligand-bound receptor crystal structures derived from X-ray diffraction data can bias conformations such that the ground or fully active state is observed but the spectrum of graded or partial activity states are not, solution NMR spectroscopy studies are capable of detecting graded activity conformational states (Boehr et al. 2009, Kar et al. 2010, Casiraghi et al. 2019). Furthermore, to our knowledge, a direct quantitative assessment has yet to be reported on the relationship between graded ligand potency, efficacy, and receptor conformation as predicted by the two-state model of receptor activation (Leff 1995). Does ligand potency correlate with functional efficacy within a structurally related series of ligands, or among a structurally diverse set of ligands?

Here we present a quantitative receptor activity-conformation analysis using two distinct sets of agonists of the nuclear receptor peroxisome proliferator-activated receptor gamma (PPARγ). We characterized a series of 10 structurally related synthetic PPARγ agonists spanning ~10,000-fold in affinity using biochemical, biophysical, and cellular assays and structural analysis using X-ray crystallography and NMR spectroscopy. Ligand potency and efficacy in this series is correlated to the degree to which the ligands shift the conformational ensemble of PPARγ towards an active state, which we detected by NMR but not in crystal structures, in a manner consistent with relationships predicted by the two-state model of receptor activation (Leff 1995) and the CTC model. However, using a larger structurally diverse set of ligands including various endogenous and synthetic PPARγ ligands, we found that ligand efficacy and receptor conformation are correlated independent of ligand potency. Collectively, our studies that data from quantitative structure-function approaches can predict ligand efficacy and assess theoretical models of receptor function.

## RESULTS

### Graded potency and efficacy within a structurally related series of PPARγ agonists

We assembled a series of 10 thiazolidinedione (TZD) PPARγ agonists (Fig. 1a) that include several FDA-approved antidiabetic drugs (Willson et al. 1996). All ligands within series contain the conserved TZD head group connected by a linker to a central aromatic moiety and a variable tail group. The linker in all but one of the ligands is a flexible, saturated methylene group that links the TZD head group to a central aromatic moiety; CAY10638 contains an unsaturated linker, which restricts the mobility of the TZD head group. The central aromatic moieties mostly comprise phenyl moieties with the exception of naphthalene and benzothiophene moieties in Netoglitazone and Edaglitazone, respectively. In contrast to these relatively conservative changes near the TZD head group, the series encompasses a variety of tail moieties extended from the central aromatic core.

**Figure 1.**
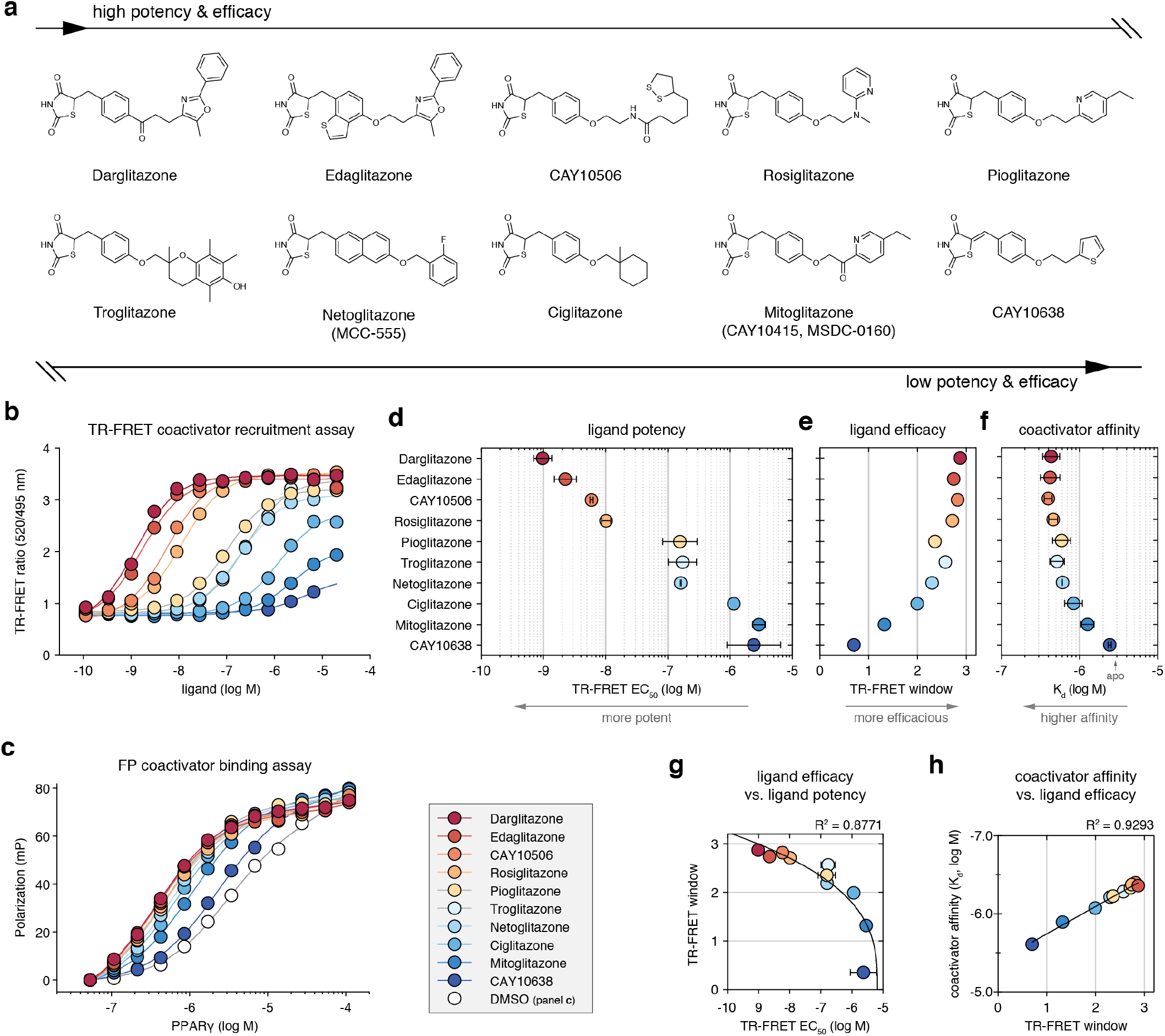
Quantitative potency and efficacy characterization of a series of thiazolidinedione (TZD) PPARγ agonists. (**a**) Chemical structures of the TZD ligands. (**b**) TR-FRET assay to determine ligand potency (EC_50_ values) and efficacy (TR-FRET window of activity) for recruitment of TRAP220 coactivator peptide to PPARγ LBD fit to a sigmoidal dose response equation; error bars, mean ± s.d. (n=3). (**c**) FP assay to determine TRAP220 coactivator peptide affinity to PPARγ LBD when bound to the ligands fit to a one site — total binding equation; error bars, mean ± s.e.m (n=3). (**d**) Ligand potencies from the fitted TR-FRET data; error bars, mean ± s.d. of two replicate experiments. (**e**) Ligand efficacies from the fitted TR-FRET data of one experimental replicate; there is experiment-to-experiment variation in the window magnitude but not the overall trends for the TZD series. (**f**) TRAP220 peptide affinities from the fitted FP data; error bars, fitted error. The affinity for apo-protein is indicated on the x-axis by a gray arrow. (**g**) Plot of ligand efficacy from (**e**) vs. ligand potency from (**d**) fit with a cubic polynomial equation. (**h**) Plot of TRAP220 peptide affinity from (**f**) vs. ligand efficacy from (**e**) fit with a linear regression equation.

Agonists increase PPARγ-mediated transcription by enhancing binding of transcriptional coactivator proteins, such as TRAP220, also known as MED1 or DRIP205 (Ge et al. 2002). We assessed the activities of the TZD series in two quantitative biochemical assays. We used a time-resolved fluorescence resonance energy transfer (TR-FRET) biochemical assay that measures the ligand-dependent change in the interaction between the PPARγ LBD and a peptide derived from the TRAP220 coactivator (Fig. 1b) containing an “LXXLL” nuclear receptor interaction motif (Savkur et al. 2004). In the TR-FRET coactivator recruitment assay, differences in the overall assay window (efficacy), which relates to the relative degree of TRAP220 peptide recruitment, are indicative of ligand-dependent differences in the binding affinity of the TRAP220 coactivator peptide to the PPARγ LBD, which we probed directly using a fluorescence polarization assay (Fig. 1c).

In the TR-FRET coactivator recruitment assay, the TZD series spans nearly 10,000-fold in potency (EC_50_), or five orders of magnitude (Fig. 1d), and showed decreasing coactivator peptide recruitment efficacy (Fig. 1e) as ligand potency decreases (Fig. 1g). Coactivator binding affinity was also decreased when PPARγ was bound to less potent ligands (Fig. 1f). Consistent with the correlation between TR-FRET recruitment efficacy and coactivator affinity there is a correlation between coactivator affinity and ligand efficacy (Fig. 1h). This correlated pattern of decreased ligand potency and efficacy is consistent with the allosteric principles of the two-state model of receptor activation (Leff 1995), which describes the actions of full and partial, or less efficacious, agonists where decreasing or graded potency within a ligand series is associated with graded efficacy (Black et al. 1985, Kenakin 2017).

### Correlation of TZD affinity, receptor stability, and cellular transcription

We determined ligand binding affinities for the TZD series using a TR-FRET assay that measures the displacement of a fluorescent tracer ligand (Fig. 2a). The TZD series spans an affinity (K_i_) range of five orders of magnitude (Fig. 2b) and is highly correlated to biochemical potency in the TR-FRET assay (Fig. 2e). Ligand binding affinity is generally correlated to a proportional increase in the thermal stability of the ligand-bound receptor (Cimmperman et al. 2008). Using differential scanning calorimetry, we determined the unfolding temperature of the PPARγ LBD when bound to each ligand in the TZD series (Fig. 2c) and found a linear correlation between ligand binding affinity and receptor stability (Fig. 2f). We also assessed the TZD series in cell-based transcriptional luciferase reporter assay that directly reports on the transcriptional activity of the PPARγ LBD (Fig. 2d) and is highly sensitive to graded PPARγ agonism (Hughes et al. 2012). The cellular transcription profile of the TZD series is similar to the quantitative biochemical TR-FRET coactivator recruitment profiles. Due to cellular toxicity of the ligands at higher concentrations, we were unable to determine cellular potency values for some TZDs due to non-saturating cellular response profiles. However, there is a correlation between the maximal luciferase value derived a fitted of the transcriptional reporter assay and TR-FRET ligand potency (Fig. 2g) and ligand affinity (Fig. 2h). Thus, ligand affinity, receptor stability, and transcription is correlated to quantitative functional potency and efficacy within the TZD series.

**Figure 2.**
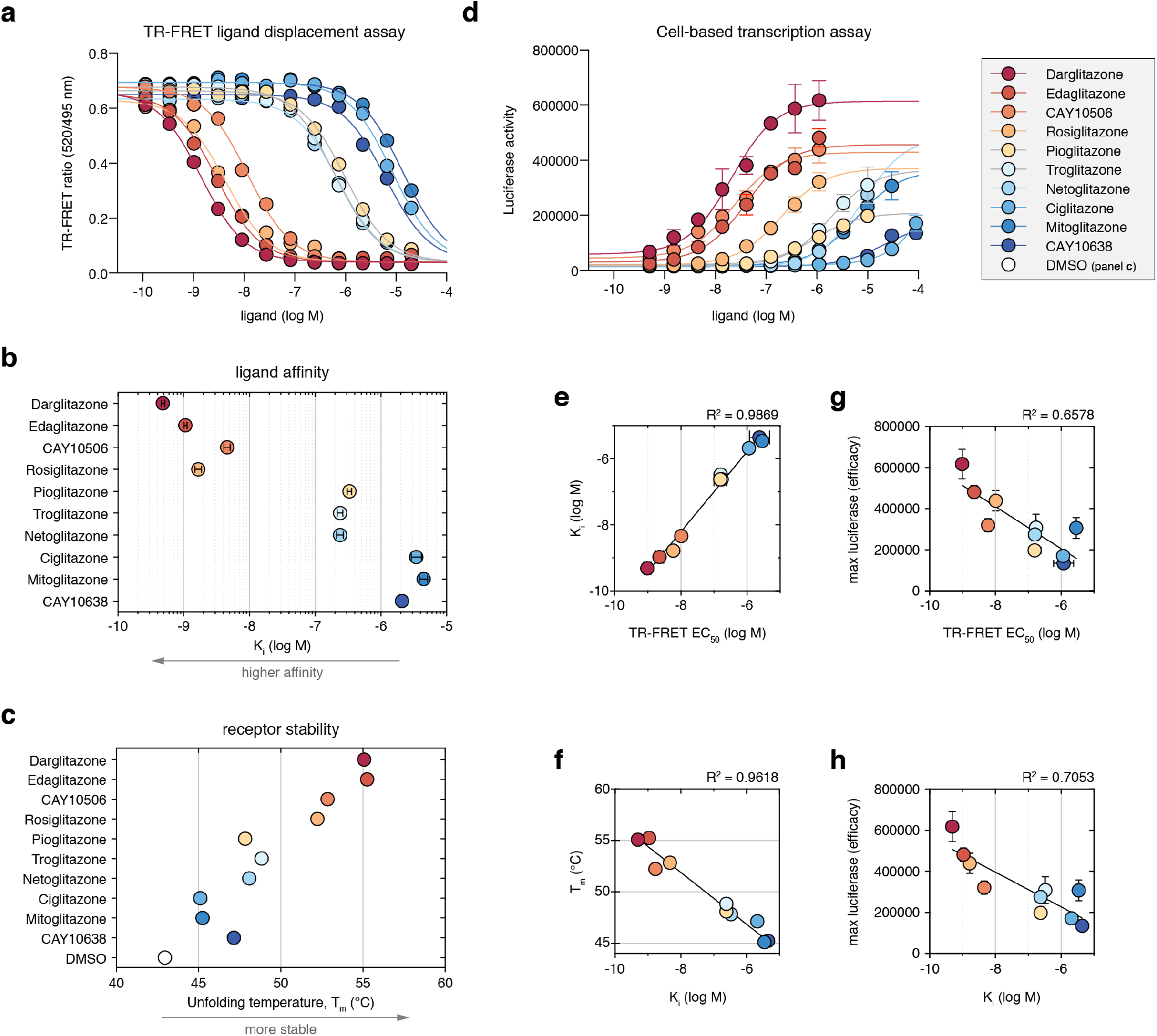
Affinity of the TZD series and effects on receptor stability and transcription. (**a**) TR-FRET assay to determine ligand affinity (K_i_ values) using a fluorescent tracer ligand fit to the Cheng-Prusoff inhibitor constant equation; error bars, mean ± s.d. (n=3). (**b**) Ligand affinities (K_i_ values) from the fitted TR-FRET data; error bars, mean ± s.d. of two replicate experiments. (**c**) PPARγ LBD thermal unfolding/melting temperatures from fitted differential scanning calorimetry (DSC) data. (**d**) Cell based luciferase assay reporting on transcription of the PPARγ LBD fit to a sigmoidal dose response equation; error bars, mean ± s.d. (n=4). (**e**) Plot of ligand affinity from (**b**) vs. ligand potency from the TR-FRET coactivator recruitment assay from (Fig. 1d) fit with a linear regression equation. (**f**) Plot of receptor stability from (**c**) vs. ligand affinity from (**b**) fit with a linear regression equation. (**g,h**) Plot of transcriptional window of efficacy via maximum luciferase activity from the highest ligand dose in (**d**) vs. (**g**) ligand potency in the TR-FRET coactivator recruitment assay from (Fig. 1d) and (**h**) ligand affinity in the TR-FRET ligand displacement assay from (**b**) fit with a linear regression equation.

### Crystal structures provide some insight into TZD affinity but not efficacy

To gain insight into the structural basis for the varied TZD affinity, potency, and functional efficacy, we compared crystal structures of PPARγ LBD bound to each of the TZDs. We solved crystal structures of six different complexes (**Supplementary Table S1**) of PPARγ LBD bound to Darglitazone (1.95 Å resolution), CAY10506 (2.45 Å resolution), Troglitazone (3.10 Å resolution), Ciglitazone (2.78 Å resolution), Mitoglitazone (2.52 Å resolution), and CAY10638 (2.15 Å resolution). We compared our structures to four previously solved PPARγ LBD crystal structures bound to Edaglitazone (PDB: 5UGM; 2.1 Å resolution) (Shang et al. 2018), Rosiglitazone (PDB: 4EMA; 2.55 Å resolution) (Liberato et al. 2012), Pioglitazone (PDB: 5Y2O; 1.80 Å resolution) (Lee et al. 2017), and Netoglitazone/MCC-155 (PDB: 3B0Q; 2.10 Å resolution). This enabled a complete X-ray crystallographic analysis of the TZD series (Fig. 3a). In all of the structures, PPARγ LBD crystallized with two molecules configured as a homodimer in the asymmetric unit, although in solution is the PPARγ LBD is monomeric (Bernardes et al. 2012). Similar to other PPARγ LBD crystal structures solved in the absence of coregulator peptide, in chain A the critical switch element for activation of PPARγ transcription, helix 12, adopts an active conformation. The conformation of helix 12 in chain B is atypical and influenced by a crystallization artifact; it binds to the AF-2 surface of an adjacent molecule (chain A’), which likely stabilizes the active conformation of helix 12 in chain A.

**Figure 3.**
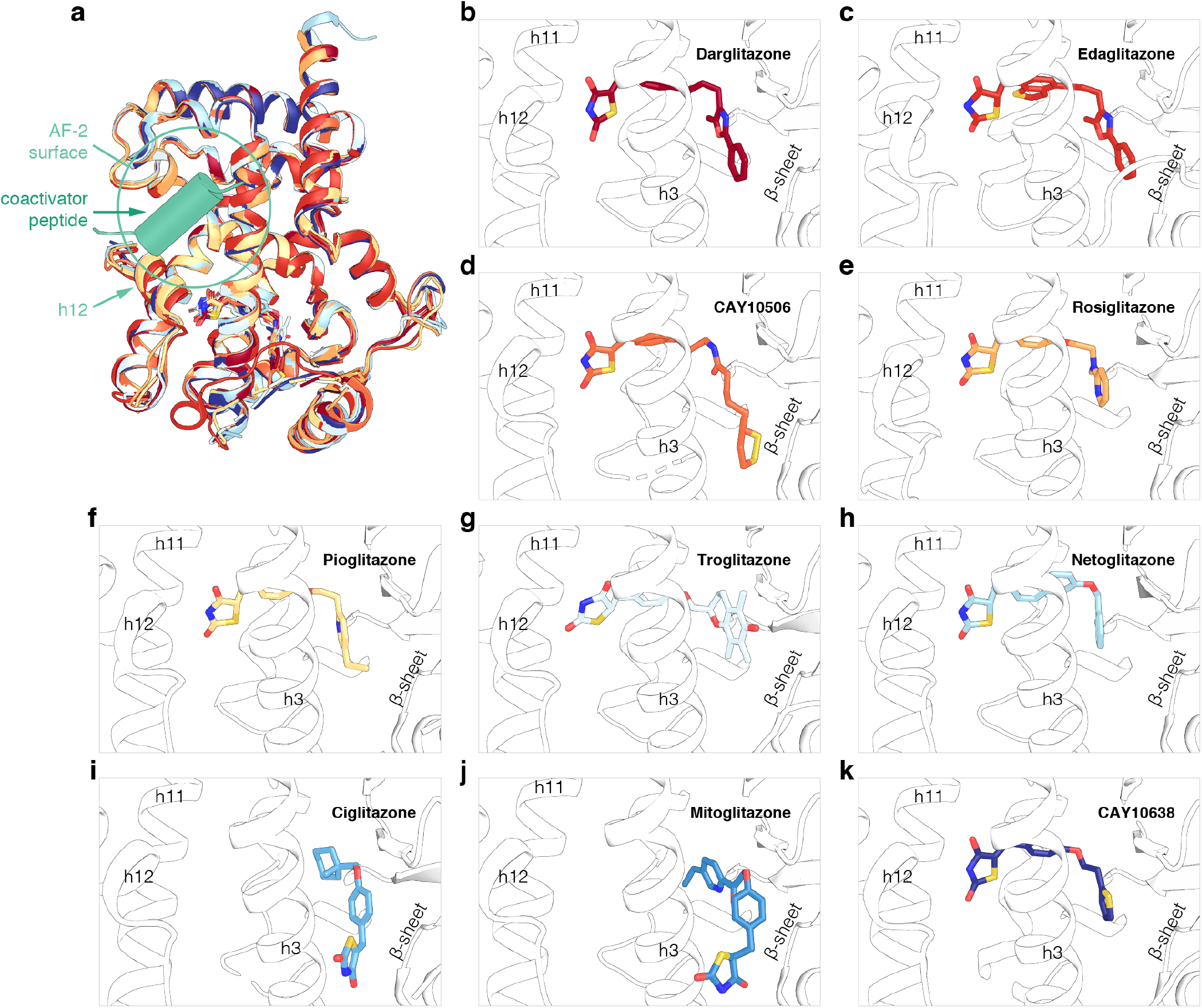
X-ray crystallography analysis of the TZD series. (**a**) Structural overlay of the ten TZD-bound PPARγ LBD structures (chain A). (**b–k**) Ligand binding poses of the TZD series; (**b**) Darglitazone (PDB 6DGL), (**c**) Edaglitazone (PDB 5UGM), (**d**) CAY10506 (PDB 6DGQ), (**e**) Rosiglitazone (PDB 4EMA), (**f**) Pioglitazone (PDB 5Y2O), (**g**) Troglitazone (PDB 6DGO), (**h**) Netoglitazone (PDB 3B0Q), (**i**) Ciglitazone (PDB 6O68), (**j**) Mitoglitazone (PDB 6O67), and (**k**) CAY10638 (PDB 6DGR).

All ligands in the TZD series show clear density in the chain A molecule (Fig. 3b–k; **Supplementary Figs. S1 and S2**). For 8 of the 10 ligands (all except Ciglitazone and Mitoglitazone), the TZD head group associates near helix 12, a region of the ligand-binding pocket called the helix 12 subpocket herein, and forms hydrogen bond contacts with the side chains of up to four nearby residues Ser289, His323, His449, and Tyr473. Among these ligands, there is a trend in the ligand binding poses whereby the higher affinity, most potent ligands display weaker polar interactions with the side chains of Cys285 and/or Gln286 and contain longer tail groups that wrap around helix 3 and form water-mediated polar interactions with residues in the β-sheet. In contrast to these 8 ligands that adopt “canonical” binding modes, Ciglitazone and Mitoglitazone, two of the least potent ligands in the series, show alternate binding modes in the chain A active conformation where their TZD head groups associate near the flexible, solvent accessible Ω-loop and their relatively shorter extended tail groups insert into the orthosteric pocket along side the β-sheet surface. Ligand binding to this alternate site has been observed in several other structural studies (Hughes et al. 2014, Bae et al. 2016, Hughes et al. 2016, Brust et al. 2017, Jang et al. 2017, Laghezza et al. 2018). The alternate binding modes of Ciglitazone and Mitoglitazone may originate from their lower affinity, and in the case of Ciglitazone, a lack of atoms in its shorter tail group capable of forming water-mediated polar interactions with residues in the β-sheet (Mosure et al. 2019). Although the significance of these alternate TZD binding modes is not yet clear, it is possible they represent an initial encounter complex binding mode before transitioning to the orthosteric ligand-binding pocket.

Whereas the structural contributions to the graded ligand binding affinity within the TZD series may be apparent from the crystallized ligand binding modes (Fig. 3b–k), the structures do not explain the graded functional efficacy of the TZD series. In principle the differences in graded efficacy should manifest in conformational differences in the structural elements that comprise the activation function-2 (AF-2) coregulator interaction surface. However, there are no obvious structural changes in the AF-2 surface among the crystal structures for this TZD series (Fig. 3a). This is likely due to the aforementioned helix 12/AF-2 surface crystal contacts, although another contribution could be that the crystallized conformations do not fully represent the conformational ensemble of PPARγ in solution. Namely, the crystallized conformations we captured may represent a low energy minima of a broader energy landscape that are not sensitive to differences caused by ligand binding in X-ray data collected under cryogenic temperatures (Fraser et al. 2009, Fraser et al. 2011, Tyka et al. 2011, Keedy et al. 2015).

### NMR reveals graded TZD potency and efficacy is correlated to receptor active state

Previous NMR studies have shown that the ligand-binding pocket and helix 12 of apo-PPARγ LBD is dynamic and switches between two or more conformations on the microsecond-millisecond (μs-ms) timescale, also known as the intermediate exchange NMR timescale, resulting in the appearance of approximately half of the expected NMR peaks (Johnson et al. 2000, Hughes et al. 2012). NMR peaks missing in the apo-form, which include residues within ligand-binding pocket and the AF-2 coregulator interaction surface (helix 3, helix 4/5, and helix 12), are stabilized upon binding potent full agonists that robustly activate PPARγ but remain absent, or persists, upon binding potent non-TZD partial agonists that weakly activate PPARγ (Johnson et al. 2000, Berger et al. 2003, Hughes et al. 2012, Marciano et al. 2015). This suggests that μs-ms timescale dynamics in the ligand-binding pocket and AF-2 surface that persist from apo-form to the ligand-bound form cause dynamical dysfunction (Mauldin et al. 2009, Peng 2009) whereby the interaction of coactivator proteins with PPARγ bound to potent partial agonists is not favored compared to potent full agonists (Kojetin et al. 2013).

The two-state model of receptor activation (Leff 1995) relates the rate of transformation of the receptor conformational ensemble from a ground state to the ligand-bound active state (Kenakin 2014, Kenakin 2017). Solution NMR spectroscopy is a powerful approach to study dynamic allosteric properties in proteins (Grutsch et al. 2016). We therefore envisioned that NMR may detect a shift in the PPARγ LBD conformational ensemble towards the activate state for the TZD series in a manner consistent with the two-state model.

We performed differential NMR analysis comparing two-dimensional [^1^H,^15^N]-TROSY-HSQC NMR spectra of ^15^N-labeled PPARγ LBD bound to each ligand within the TZD series to the apo-form under the same ligand vehicle DMSO condition. Ligands within the series contain different extended side chain moieties, including various aromatic groups that interact with the β-sheet region and wrap around helix 3 resulting in ring current effects that impart unpredictable nonlinear NMR peak shifting for residues within this region of the pocket. NMR peak shifting is also observed for residues near the ligand entry/exit site, likely due in part to slight differences in the relative binding mode of the ligands. Despite these general binding mode-induced NMR peak shifting, two general NMR observable conformational trends are apparent for the TZD series.

We found that the less potent ligands within the series that caused partial or graded functional efficacy resulted in NMR peak line broadening (Fig. 4a, **Supplementary Figure S4**), indicative of μs-ms timescale dynamics, that persist from the apo-form for residues within the ligand entry/exit site, ligand-binding pocket, and structural elements comprising the AF-2 coregulator interaction surface (Fig. 4c). The structural regions with persistent μs-ms timescale dynamics when bound to the less potent TZDs comprise similar structural regions with persistent μs-ms timescale dynamics when bound to potent non-TZD partial agonists. In the case of the less potent TZD ligands, the persistent μs-ms timescale dynamics likely originates from relatively fast ligand off-exchange compared to the more potent TZDs (Carroll et al. 2012). However, for the potent non-TZD partial agonists, the persistent μs-ms timescale dynamics likely originates from a different mechanism not related to ligand off-exchange; these ligands were developed to retain high affinity but lack a TZD-comparable head group capable of forming hydrogen bond contacts to residues within the pocket and on helix 12 that stabilize the AF-2 surface. Thus, when bound to PPARγ at high affinity, the non-TZD partial agonists enable the dysfunctional μs-ms timescale dynamics present in the apo-form to persist.

**Figure 4.**
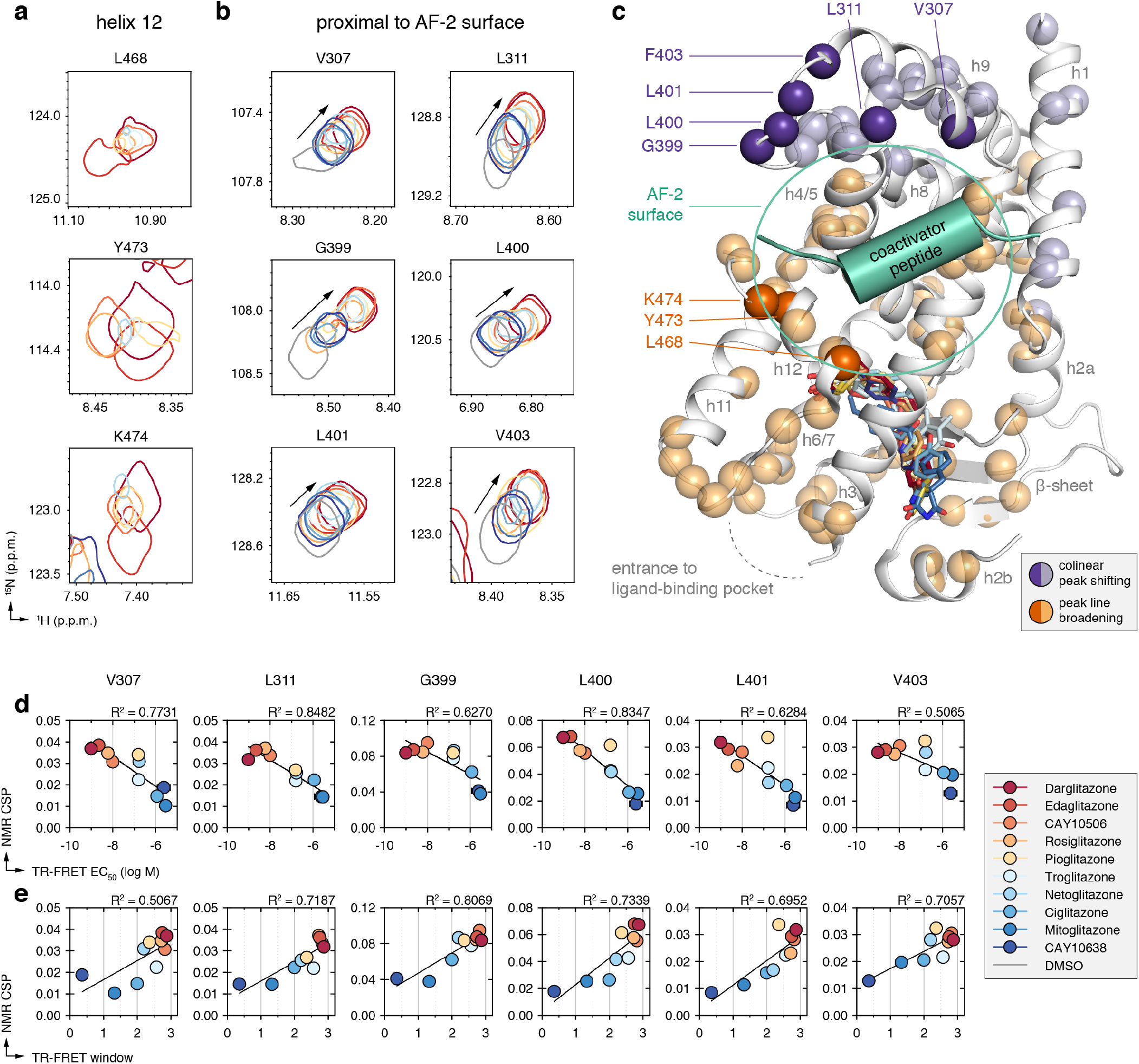
NMR-detected changes in the PPARγ LBD conformational ensemble correlate with graded ligand potency and efficacy. (**a,b**) Snapshots of [^1^H,^15^N]-TROSY-HSQC NMR spectra of ^15^N-labeled PPARγ LBD that show NMR changes as a function of graded potency and efficacy, including residues (**a**) in helix 12 that show NMR peak line broadening or (**b**) proximal to the AF-2 coactivator interaction surface that show co-linear shifting. (**c**) Analysis of all well-dispersed NMR peaks with reasonable peak separation that could be faithfully analyzed shows widespread correlations throughout the PPARγ LBD that group into two surfaces sensitive either to NMR peak line broadening or co-linear shifting. Dark orange and dark purple spheres correspond to residues shown in (**a,b**); light orange and light purple spheres correspond to other residues that also show line broadening (**Supplementary Figure S4**) or co-linear shifting (**Supplementary Figure S5**) in the TZD series. (**d,e**) Plot of NMR chemical shift perturbation for each liganded state shown in (**b**) relative to the apo-protein state vs. (**d**) ligand potency from (Fig. 1d) and (**e**) ligand efficacy in the TR-FRET coactivator recruitment assay from from (Fig. 1e) fit with a linear regression equation.

In contrast to these NMR line broadening changes, we observed co-linear NMR chemical shift perturbations for residues distal from the ligand-binding pocket but proximal to the AF-2 surface (Fig. 4b, **Supplementary Figure S5**), which includes residues that form a surface connected by helix 1, helix 4/5, helix 8, helix 8-9 loop, and helix 9 (Fig. 4c). In many cases, one NMR peak is visible when bound to the most potent ligands or least potent ligands, whereas ligand with intermediate potency show a combination of shifting, line broadening, and two receptor populations. The coincident shifting and broadening suggests conversion on the fast (peak shifting) to intermediate (peak broadening) NMR timescale. The two receptor populations observed for the intermediate potency ligands indicate there is exchange between two ligand-bound receptor populations. One likely contribution to this phenomenon is the racemic nature of the TZD head group, which isomerize between *(R)*- and *(S)*-conformers, each of which have different affinities for binding (Mosure et al. 2019). For example, *(S)*-Rosiglitazone displays higher affinity than *(R)*-Rosiglitazone (30 nM vs. 2 μM, respectively). It is possible that as the overall racemic affinity is reduced, the difference in affinity between isomers becomes less significant allowing both to bind. In this scenario, the different TZD head group isomers will interact differently with the receptor, which could in principle cause the peak doubling observed that correspond to two receptor populations. Notwithstanding the isomer contributions, the degree to which the NMR peak shifts from a ground state conformation (apo-form) towards a fully “active” conformation is correlated to ligand potency (Fig. 4d). When considered with our other quantitative analyses, these graded activity conformations are also correlated to ligand efficacy (Figs. 1 and 2) in a manner consistent with the two-state model of receptor activation (Leff 1995).

### Ligand efficacy is correlated to receptor active state within a larger set of structurally diverse ligands

We wondered whether there may be a correlation between ligand efficacy and receptor conformation within a different set of structurally diverse ligands. We assembled 18 natural/ endogenous and synthetic PPARγ ligands not previously reported to span a range of graded transcriptional activation efficacy (**Supplementary Fig. S3**). Unlike the TZD series, which share a common head group that interacts with the helix 12 subpocket, this diverse ligand set contains different types of head groups and chemotypes, some capable of hydrogen bonding to the helix 12 subpocket and others lacking such moieties that function primarily as partial agonists. Given the diversity of this ligand set, some of which were optimized for high affinity binding with low transcriptional efficacy by incorporating different types of head groups, we initiated these studies understanding that ligand potency will not be correlated to efficacy as we observed in the more structurally conserved TZD series.

We compared two-dimensional [^1^H,^15^N]-TROSY-HSQC NMR spectra of ^15^N-labeled PPARγ LBD bound to each ligand within the structurally diverse set. Due to the diversity of chemotypes present in this ligand set, there are larger differences in the NMR spectra compared to the TZD series. However, several residues with well-resolved NMR peaks that showed co-linear shifting in the TZD series also showed co-linear shifting in this structurally diverse ligand set (Fig. 5a), indicating a correlation between ligand efficacy and receptor conformation. Compared to the TZD series NMR data, the majority of natural/endogenous PPARγ ligands show shifted NMR peak positions indicating they are partial agonists, which is consistent with the idea that PPARγ displays basal transcriptional activity and can be further activated by synthetic agonists (Shang et al. 2018). To assess this more directly, we performed our TR-FRET coactivator recruitment assay using a single ligand concentration (5 μM) to assess ligand efficacy (Fig. 5b). In contrast to the NMR experiments where the receptor concentration is relatively high (200 μM) and ligands added stoichiometrically result in ~100% bound occupancy, in the TR-FRET assay the receptor concentration is low (4 nM) and ligands titrated increase binding of the peptide (efficacy) in proportion to their respective binding affinities. Because high concentrations of excess ligand can result in compound precipitation or colloidal aggregate formation, we limited this analysis to moderately high affinity synthetic PPARγ ligands with affinities better than ~1–2 μM (**Supplementary Fig. S6**) that would show appreciable complex formation under the TR-FRET assay conditions. For this ligand subset, we observed a correlation between TR-FRET coactivator recruitment efficacy and co-linear NMR peak shifting (Fig. 5c). However, for these ligands there is a poor correlation between ligand affinity and efficacy (Fig. 5d). For example, MRL24 is a partial agonist with a low TR-FRET efficacy window value similar to GQ-16; however, MRL20 is much more potent than GQ-16 and is also more potent than the super agonist GW1929, which has the highest TR-FRET efficacy value. Thus, for the structurally diverse ligand set, these data show that the receptor conformation shifts from a ground state conformation (apo-form) towards an active conformation in a manner that is correlated to ligand efficacy but independent of ligand affinity.

**Figure 5.**
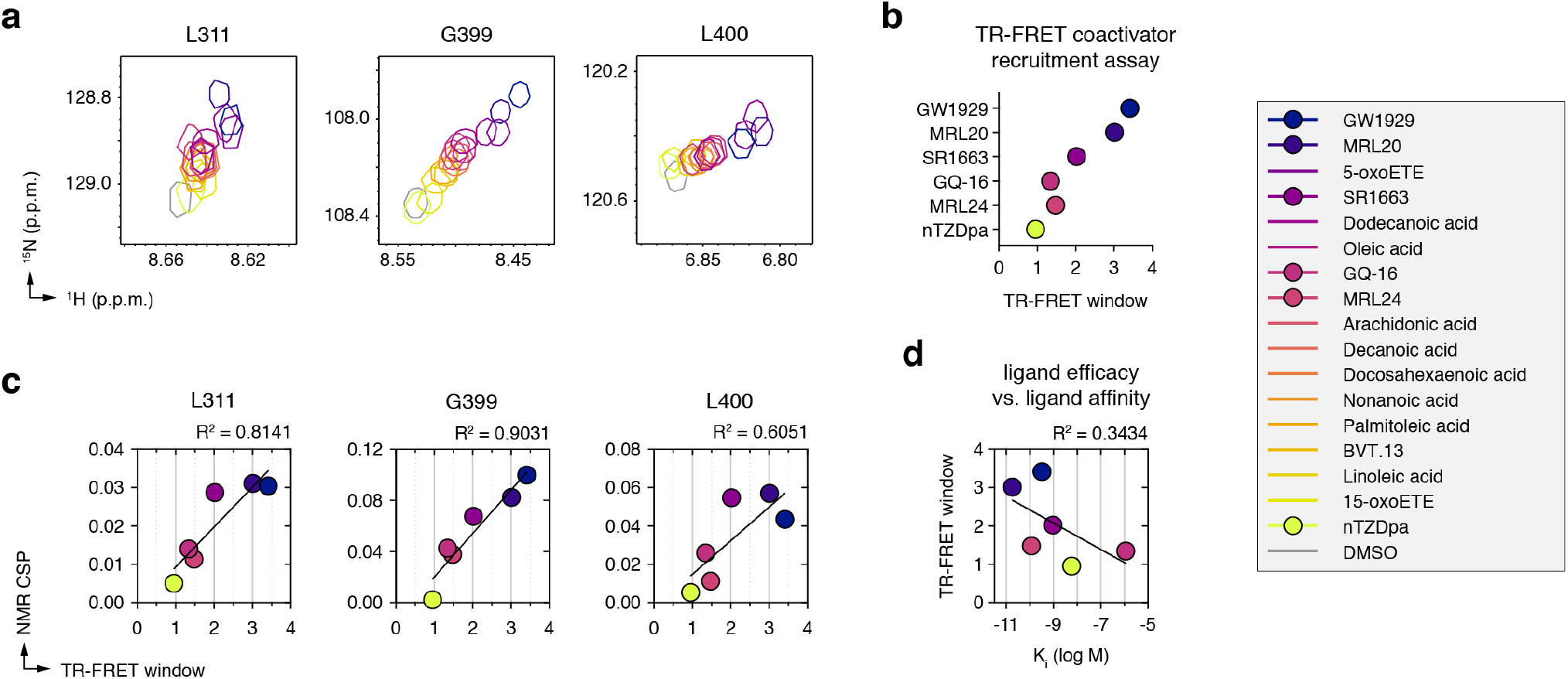
NMR-detected correlations PPARγ LBD active state and efficacy of a structurally diverse ligand set. (**a**) Snapshots of [^1^H,^15^N]-TROSY-HSQC NMR spectra of ^15^N-labeled PPARγ LBD that show co-linear NMR peak shifting for residues proximal to the AF-2 coactivator interaction surface. (**b**) TR-FRET assay to determine ligand efficacy (TR-FRET window of activity) for recruitment of TRAP220 coactivator peptide to PPARγ LBD at a single concentration of ligand (5 μM) fit to a sigmoidal dose response equation; error bars, mean ± s.e.m (n=3). In the legend, synthetic ligands with a filled circle correspond to ligands with affinities (K_i_ values) better than ~1–2 μM that were included in the TR-FRET assay. (**c**) Plot of NMR chemical shift perturbation for each liganded state relative to the apo-protein state vs. ligand efficacy in the TR-FRET coactivator recruitment assay from (**b**) fit with a linear regression equation. (**d**) Synthetic ligands included in the TR-FRET and NMR cross-correlation analysis show poor correlation to ligand efficacy.

## DISCUSSION

The objective of receptor theory is to understand and predict the relationship between ligand potency and functional efficacy. Theoretical receptor models, which are used to extract quantitative information from functional activity studies on GPCRs, ion channels, and enzyme-linked receptors, have evolved over the years to account for new findings in membrane receptor function: the realization that GPCRs can affect multiple signaling pathways and therefore have multiple distinct active states, the discovery of orthsoteric vs. allosteric ligand binding sites, among others. Computer simulations are heavily used in the development and assessment of theoretical receptor models by varying mathematical parameters to determine relationships between ligand potency, functional efficacy, and other model system parameters. However, the manner in which the ligand affects receptor conformation in these theoretical studies is a conceptual “black box”—there is no knowledge or description of how ligand binding pose or chemical modifications influence ligand binding affinity, receptor conformation, and receptor functional activity. To gain insight into this relationship, here we studied the influence of chemical modifications within a structurally related ligand series (TZDs) and a structural distinct ligand set using quantitative experimental approaches that enable assessment of ligand potency, graded functional efficacy or activity, and receptor conformation.

To our knowledge, theoretical receptor models such as the two-state model of receptor activation have not been utilized or considered in studies of nuclear receptor transcription factors, but our work here shows they are relevant. We used several quantitative functional assays that report on various ligand-responsive biophysical and functional endpoints related to nuclear receptor function including direct physical readouts on the PPARγ LBD (ligand displacement and thermal stability assays) as well as coactivator affinity for and recruitment to the PPARγ LBD (TR-FRET and FP assays), the latter of which has a direct relation to cellular transcription (luciferase assay). As predicted by the two-state model, we observed correlations between the various functional activity readouts for PPARγ agonists within the structurally related TZD series. To relate the graded activities of the TZD series to receptor conformation, we performed structural analysis of ligand-bound PPARγ LBD complexes using X-ray crystallography and NMR spectroscopy. We did not observe any apparent graded activity conformations in crystal structures of PPARγ LBD bound to the TZD series. However, NMR detected a graded shift in the conformational ensemble of the PPARγ LBD towards an active conformation correlated to the degree of ligand potency and efficacy, and therefore also correlated to the other functional activity readouts (ligand affinity, receptor stability, coactivator affinity)—all consistent with underpinnings of allostery in the two-state model of receptor activation. We were unable to directly measure the binding kinetics of the TZD series due to apparent fast binding kinetics in surface plasmon resonance (SPR) experiments, suggesting a two-state binding mechanism. However, assuming that all of the TZDs display similar association rate constants, the notion that all of the functional efficacy and conformational data correlate to ligand affinity indicates that ligand residence times (i.e., ligand dissociation kinetic rate constants) are directly involved in the functional activities of the TZD series.

In contrast to our results with the structurally related TZD agonist series, we found that the predicted relationship in the two-state model between ligand potency and efficacy breaks down for structurally diverse endogenous and synthetic PPARγ agonists. That is, whereas ligand binding affinity or potency can predict functional efficacy within the structurally related agonists, ligand affinity or potency does not predict the functional efficacy of structurally distinct agonists. On the structural level, this can be rationalized because different ligand scaffolds, or even small chemical modifications, result in different chemical bonding patterns between ligand and receptor that can influence affinity distinct from functional efficacy. For the TZD series, these ligands have a conserved chemical group (TZD) capable of hydrogen bonding to residues within the helix 12 subpocket, in particular Y473 on helix 12, which stabilizes the AF-2 surface into an active conformation. The other chemical changes within the TZD series, in particular the varied length of tail groups that contribute to ligand affinity, are all structurally distal from the AF-2 surface. In contrast, the structurally distinct ligands possess different types of head groups that localize near the helix 12 subpocket, some capable of robustly stabilizing the AF-2 surface and others that are not. Furthermore, within this diverse ligand set there are many other chemical variations in the tail groups that distinctly contribute to affinity. Many of the endogenous agonists possess relatively simple aliphatic chain tail groups whereas the synthetic agonists contain chemically diverse tail groups; all of these factors have a significant influence on ligand binding affinity independent of the ability of the head group to stabilize the AF-2 surface. Although these diverse ligands contradicted the potency-efficacy correlation predicted by the two-state model, our NMR analysis revealed a correlation between efficacy and graded activity conformation state for all ligands, independent of chemical structure.

Related to our work here on nuclear receptors, NMR studies have been used to define how ligand binding influences the conformational ensemble of GPCRs in ways not detectable in ligand-bound crystal structures. One-dimensional ^19^F NMR studies of the GPCRs β_2_-adrenergic receptor (β_2_AR) and adenosine A_2_ receptor (A_2_AR) tagged with a ^19^F NMR reporter molecule attached to two cysteine residues revealed that pharmacologically distinct classes of ligands or nanobodies differentially stabilize the receptor conformational ensemble into G-protein inactive and active states (Liu et al. 2012, Kim et al. 2013, Manglik et al. 2015, Staus et al. 2016, Ye et al. 2016, Prosser et al. 2017, Susac et al. 2018). We similarly used ^19^F NMR to show that pharmacologically distinct PPARγ ligands differentially affect the conformation of the AF-2 helix using a ^19^F NMR reporter molecule attached to helix 12, and the conformations we observed are predictive of ligand efficacy (Chrisman et al. 2018). Furthermore, using selective labeling approaches and two-dimensional NMR, other work has revealed the existence of ligand-bound conformational states that exist in the continuum between the inactive/ground state and fully active conformations previously captured in crystal structures (Bokoch et al. 2010, Kofuku et al. 2012, Nygaard et al. 2013, Kofuku et al. 2014, Okude et al. 2015, Sounier et al. 2015, Isogai et al. 2016, Clark et al. 2017, Solt et al. 2017, Eddy et al. 2018). Among these, most related to our study here is an NMR analysis of backbone amide groups of valine residues in β_1_-adrenergic receptor (β_1_AR) bound to structurally diverse agonists and antagonists (Isogai et al. 2016), which observed a heterogenous conformational response to the ligands. Within the structurally diverse β_1_AR ligand set, there did not seem to be any relationship between ligand affinity and functional efficacy. The conformation of several residues within the extracellular ligand binding pocket were differentially correlated to ligand affinity and ligand chemical composition, whereas the conformation of a different group of residues on the intracellular side of transmembrane helix 5 (TM5) correlated to G protein ligand efficacy independent of ligand affinity. This is similar to our finding here that structurally distinct PPARγ agonists stabilize an active conformation to a degree correlated to their functional efficacy independent of ligand potency.

Our findings raise a question as to whether theoretical receptor models can be updated to better predict the relationship between ligand potency with functional efficacy and receptor conformation. This would not be a trivial task as it would require knowledge of ligand binding poses and a precise understanding of how small chemical modifications affect ligand affinity and stabilization of functional surfaces— two distinct but important consequences of ligand-receptor interactions. In structure-activity relationship (SAR) studies, it is difficult if not impossible to predict how small chemical changes in ligand composition affects potency and efficacy (Fujioka et al. 2012, Dosa et al. 2016). There are also structural differences in how different ligand-binding proteins bind synthetic ligands that could influence potency-efficacy relationships. In nuclear receptors, residues within the ligand-binding pocket that contact the ligand are connected to nearby structural elements in the AF-2 coregulator binding surface. In contrast, the extracellular ligand-binding pocket of GPCRs is structurally distant from the intracellular effector protein binding surface. It is therefore possible that small chemical differences in nuclear receptor ligands, which could impact structural elements that are closer to nuclear receptor functional surfaces could impact affinity-function relationships differently than membrane receptors. Nonetheless, our studies here on the PPARγ nuclear receptor as well as the aforementioned studies on GPCRs show that NMR analysis can predict ligand efficacy and provide novel structural insight into the influence of ligands on activity-related receptor conformational ensembles.

## ACKNOWLEDGEMENTS

We thank Sarah Mosure and Paola Munoz-Tello for helpful discussions. This work was supported in part by National Institutes of Health (NIH) grants R01DK101871 (DJK) and F32DK108442 (RB).

## AUTHOR CONTRIBUTIONS

J.S. and D.J.K conceived and designed the research. J.S. expressed and purified protein; solved the crystal structures; and performed and analyzed biochemical assays and NMR experiments. R.B. performed the cellular assays. P.R.G. and T.M.K. provided synthetic ligands that were not commercially available. D.J.K. analyzed data, supervised the research, and wrote the manuscript with J.S. and input from all authors.

## METHODS

### Materials and reagents

The TZD ligand series includes Ciglitazone, Darglitazone, Edaglitazone, Pioglitazone, Rosiglitazone, Troglitazone, Netoglitazone (MCC-555), Mitoglitazone (CAY10415, MSDC-0160), CAY10506, and CAY10638. The structurally diverse ligand set includes natural/endogenous PPARγ ligands (arachidonic acid, decanoic acid, docosahexaenoic acid, dodecanoic acid, linoleic acid, nonanoic acid, oleic acid, palmitoleic acid) and synthetic ligands (BVT.13, GQ-16, GW1929, MRL20, MRL24, nTZDpa, SR1663). All ligands except MRL20 and SR1663 were obtained from commercial sources, including BioVision (MRL24), Cayman Chemical, Sigma-Aldrich, and Tocris Bioscience; MRL20 (Bruning et al. 2007, Hughes et al. 2012, Hughes et al. 2014, Brust et al. 2017) and SR1663 (Marciano et al. 2015) were synthesized and characterized previously. A peptide containing an LXXLL nuclear receptor interaction motif from TRAP220/MED1/ DRIP205 (residues 638–656; NTKNHPMLMNLLKDNPAQD) was synthesized by LifeTein with an N-terminal FITC label with a six-carbon linker (Ahx) and an amidated C-terminus for stability.

### Protein expression and purification

Human PPARγ ligand binding domain (LBD; residues 203–477, isoform 1 numbering) was expressed in *E. coli* BL21(DE3) cells using autoinduction ZY media, or M9 minimal media supplemented with ^15^N-labeled ammonium chloride, as a Tobacco Etch Virus (TEV)-cleavable N-terminal His-tagged (6×-His) fusion protein using a pET46 Ek/LIC vector (Novagen) and purified using Ni-NTA affinity chromatography and gel filtration chromatography as previously described (Hughes et al. 2012). Purified protein was concentrated to 10 mg/mL in a buffer consisting of 20 mM potassium phosphate (pH 7.4), 50 mM potassium chloride, 5 mM TCEP, and 0.5 mM EDTA and verified by SDS-PAGE as >95% pure.

### TR-FRET ligand displacement and coregulator recruitment assays

Time-resolved fluorescence resonance energy transfer (TR-FRET) assays were performed in black 384-well plates (Greiner) with 23 μL final well volume. For the coregulator recruitment assay, each well containing 4 nM 6×His-PPARγ LBD, 1 nM LanthaScreen Elite Tb-anti-His Antibody (ThermoFisher), and 400 TRAP220 peptide in a buffer containing 20 mM potassium phosphate (pH 7.4), 50 mM potassium chloride, 5 mM TCEP, 0.005% Tween 20. TZDs were assessed in dose response format and the last (20 μM) data point for Netoglitazone and CAY10638 was removed from data fitting due to compound precipitation; other ligands were assessed as a single concentration (5 μM). For the ligand displacement assay, each well contained 1 nM 6×His-PPARγ LBD protein, 1 nM LanthaScreen Elite Tb-anti-HIS Antibody (Thermo Fisher Scientific), and 5 nM Fluormone Pan-PPAR Green (Invitrogen) in a buffer containing 20 mM potassium phosphate (pH 8), 50 mM potassium chloride, 5 mM TCEP, and 0.005% Tween-20. Compounds stocks were prepared in DMSO via serial dilution (when applicable), added to wells in triplicate, and plates were read using BioTek Synergy Neo multimode plate reader after incubation at 25 °C for 1 h. The Tb donor was excited at 340 nm; the Tb donor emission was measured at 495 nm, and the acceptor FITC emission was measured at 520 nm. Data were plotted using GraphPad Prism as TR-FRET ratio 520 nm/495 nm vs. ligand concentration (TZD series) or at a single ligand concentration (5 μM; other ligands). Coregulator recruitment data were fit to sigmoidal dose response curve equation to obtain EC_50_ and TR-FRET window values, and ligand displacement data were fit to the one site – Fit K_i_ binding equation to obtain K_i_ values using the known binding affinity of Fluormone Pan-PPAR Green (2.8 nM; Invitrogen PV4894 product insert).

### Fluorescence polarization coregulator interaction assay

6×His-PPARγ LBD was diluted by serial dilution into a buffer containing 20 mM potassium phosphate (pH 8), 50 mM potassium chloride, 5 mM TCEP, 0.5 mM EDTA, and 0.01% Tween-20 and plated with 180 nM FITC-labeled TRAP220 peptide in black 384-well plates (Greiner) in the presence of DMSO (ligand vehicle) or ligand at a concentration equal to 10 μM over the highest protein concentration to ensure complete formation of ligand-bound protein in triplicate. The plate was incubated at 25°C for 1 hr, and fluorescence polarization was measured on a BioTek Synergy Neo multimode plate reader at 485 nm emission and 528 nm excitation wavelengths. Data were plotted using GraphPad Prism as fluorescence polarization signal in millipolarization units vs. protein concentration and fit to a one site — total binding equation.

### Differential scanning calorimetry

Samples containing PPARγ LBD (30 μM) incubated with 30 μM ligand or 0.15% DMSO (vehicle control) were degassed for 10 minutes in a buffer containing 20 mM potassium phosphate (pH 7.4), 50 mM potassium chloride, 5 mM TCEP, and 0.5 mM EDTA; 1 mL sample aliquots were transferred to duplicate wells of a 96-well deep-well plate and loaded to the autosampler module of Nano DSC (TA Instruments). Differential scanning calorimetry (DSC) data were obtained by increasing the temperature from 25–95 °C at a rate of 1 °C min^−1^ while monitoring the heat change of buffer, ligand-free protein in the presence of DMSO, and ligand-bound protein samples. Buffer scans were performed in triplicate before each protein denaturation experiment to condition the reference and sample cells. Baseline-corrected data were converted to molar heat capacity before defining a two-state sigmoidal integration baseline. The DSC peak was fitted with a two-state scaled model to determine the thermal unfolding/melting temperature (T_m_). Data were analyzed using TA Instruments DSC Analysis software and T_m_ values plotted in GraphPad Prism.

### Cellular transactivation assay

HEK293T (ATCC CRL03216) were cultured in Dulbecco’s Minimal Essential Medium (DMEM, Gibco) supplemented with 10% fetal bovine serum (FBS) and 50 units ml^−1^ of penicillin, streptomycin, and glutamine. Cells were grown to 90% confluency in T-75 flasks; from this, 4 million cells were seeded in a 10 cm cell culture dish for transfection using X-tremegene 9 (Roche) and Opti-MEM (Gibco) with Gal4-PPARγ LBD expression plasmid (4.5 μg) and a luciferase reporter plasmid containing the fix copies of the Gal4 Upstream Activation Sequence (5×UAS-luciferase) (4.5 μg). After an 18 hr incubation, cells were transferred to white 384-well cell culture plates (Thermo Fisher Scientific) at 10,000 cells/well in 20 μL total volume/well. After a 4 hr incubation, cells were treated in quadruplicate with 20 μL of either vehicle control (1.5% DMSO in DMEM media) or 2-fold serial dilution of TZDs. After a final 18 hr incubation, cells were harvested with 20 μL Britelite Plus (Perkin Elmer), and luminescence was measured on a BioTek Synergy Neo multimode plate reader. Data were plotted in GraphPad Prism as luminescence vs. ligand concentration and fit to a sigmoidal dose response curve.

### Crystallization and structure determination

For PPARγ LBD complexes with Ciglitazone, Troglitazone, Mitoglitazone, CAY10506, and CAY10638, the ligands were incubated at a 1:3 protein/ligand molar ratio in PBS overnight before being concentrated to 10 mg/ml and buffer exchange into phosphate buffer to remove DMSO. The PPARγ LBD complex with Darglitazone was obtained by soaking the ligand (1 mM in reservoir solution containing 5% DMSO) into preformed apo-PPARγ LBD crystals. Apo or protein/ligand complex crystals were obtained after 3–5 days at 22 °C by sitting-drop vapor diffusion against 50 μL of well solution using 96-well format crystallization plates. The crystallization drops contained 1 μL of protein (with or without ligand) mixed with 1 μL of reservoir solution containing 0.1 M MOPS (pH 7.6) or Tris (pH 7.6) and 0.8 M sodium citrate. All crystals were flash-frozen in liquid nitrogen before data collection. Data collection for PPARγ LBD bound to Ciglitazone, Mitoglitazone, CAY10506, and CAY10638 was carried out at ALS Beamline 5.0.2 at Berkeley Center for Structural Biology (Lawrence Berkeley National Laboratory). Data collection for the PPARγ LBD bound to Troglitazone and Darglitazone was carried out using our home source MicroMax007 HF x-ray generator equipped with the mar345 detector. Data were processed, integrated, and scaled with the programs Mosflm (Battye et al. 2011) and Scala in CCP4 (Winn et al. 2011). The structures were solved by molecular replacement using the program Phaser (McCoy et al. 2007) implemented in the PHENIX package (Adams et al. 2011) and using a previously published PPARγ LBD structure (PDB code: 1PRG) (Nolte et al. 1998) as the search model. The structure was refined using PHENIX with several cycles of interactive model rebuilding in COOT (Emsley et al. 2004).

### NMR spectroscopy

Two-dimensional [^1^H,^15^N]-TROSY HSQC NMR data of ^15^N-labeled PPARγ LBD (200 μM) were acquired at 298K on a Bruker 700 MHz NMR instrument equipped with a QCI cryoprobe in NMR buffer (50 mM potassium phosphate, 20 mM potassium chloride, 1 mM TCEP, pH 7.4, 10% D_2_O) with ligands added at 2 molar equivalents (TZD series) or 1 molar equivalent (structurally diverse ligands). Data were processed using Topspin 3.0 (Bruker Biospin) and analyzed using NMRViewJ (OneMoon Scientific, Inc.) (Johnson 2004), respectively. NMR chemical shift assignments previously reported for ligand-bound PPARγ (Hughes et al. 2012) were transferred to the spectra obtained in this study for well-resolved residues with conversed NMR peak positions to the previous ligand-bound forms using the minimum chemical shift perturbation procedure (Williamson 2013).

## REFERENCES

Adams, P. D., P. V. Afonine, G. Bunkoczi, V. B. Chen, N. Echols, J. J. Headd, L. W. Hung, S. Jain, G. J. Kapral, R. W. Grosse Kunstleve, A. J. McCoy, N. W. Moriarty, R. D. Oeffner, R. J. Read, D. C. Richardson, J. S. Richardson, T. C. Terwilliger and P. H. Zwart (2011). “The Phenix software for automated determination of macromolecular structures.” Methods 55(1): 94–106.

Bae, H., J. Y. Jang, S. S. Choi, J. J. Lee, H. Kim, A. Jo, K. J. Lee, J. H. Choi, S. W. Suh and S. B. Park (2016). “Mechanistic elucidation guided by covalent inhibitors for the development of anti-diabetic PPARgamma ligands.” Chem Sci 7(8): 5523–5529.

Battye, T. G., L. Kontogiannis, O. Johnson, H. R. Powell and A. G. Leslie (2011). “iMOSFLM: a new graphical interface for diffraction-image processing with MOSFLM.” Acta Crystallogr D Biol Crystallogr 67(Pt 4): 271–281.

Berger, J. P., A. E. Petro, K. L. Macnaul, L. J. Kelly, B. B. Zhang, K. Richards, A. Elbrecht, B. A. Johnson, G. Zhou, T. W. Doebber, C. Biswas, M. Parikh, N. Sharma, M. R. Tanen, G. M. Thompson, J. Ventre, A. D. Adams, R. Mosley, R. S. Surwit and D. E. Moller (2003). “Distinct properties and advantages of a novel peroxisome proliferator-activated protein [gamma] selective modulator.” Mol Endocrinol 17(4): 662–676.

Bernardes, A., F. A. Batista, M. de Oliveira Neto, A. C. Figueira, P. Webb, D. Saidemberg, M. S. Palma and I. Polikarpov (2012). “Low-resolution molecular models reveal the oligomeric state of the PPAR and the conformational organization of its domains in solution.” PLoS One 7(2): e31852.

Black, J. W. and P. Leff (1983). “Operational models of pharmacological agonism.” Proc R Soc Lond B Biol Sci 220(1219): 141–162.

Black, J. W., P. Leff, N. P. Shankley and J. Wood (1985). “An operational model of pharmacological agonism: the effect of E/[A] curve shape on agonist dissociation constant estimation.” Br J Pharmacol 84(2): 561–571.

Boehr, D. D., R. Nussinov and P. E. Wright (2009). “The role of dynamic conformational ensembles in biomolecular recognition.” Nat Chem Biol 5(11): 789–796.

Bokoch, M. P., Y. Zou, S. G. Rasmussen, C. W. Liu, R. Nygaard, D. M. Rosenbaum, J. J. Fung, H. J. Choi, F. S. Thian, T. S. Kobilka, J. D. Puglisi, W. I. Weis, L. Pardo, R. S. Prosser, L. Mueller and B. K. Kobilka (2010). “Ligand-specific regulation of the extracellular surface of a G-protein-coupled receptor.” Nature 463(7277): 108–112.

Bruning, J. B., M. J. Chalmers, S. Prasad, S. A. Busby, T. M. Kamenecka, Y. He, K. W. Nettles and P. R. Griffin (2007). “Partial agonists activate PPARgamma using a helix 12 independent mechanism.” Structure 15(10): 1258–1271.

Brust, R., H. Lin, J. Fuhrmann, A. Asteian, T. M. Kamenecka and D. J. Kojetin (2017). “Modification of the Orthosteric PPARgamma Covalent Antagonist Scaffold Yields an Improved Dual-Site Allosteric Inhibitor.” ACS Chem Biol 12(4): 969–978.

Carroll, M. J., R. V. Mauldin, A. V. Gromova, S. F. Singleton, E. J. Collins and A. L. Lee (2012). “Evidence for dynamics in proteins as a mechanism for ligand dissociation.” Nat Chem Biol 8(3): 246–252.

Casiraghi, M., E. Point, A. Pozza, K. Moncoq, J. L. Baneres and L. J. Catoire (2019). “NMR analysis of GPCR conformational landscapes and dynamics.” Mol Cell Endocrinol 484: 69–77.

Changeux, J. P. (2012). “Allostery and the Monod-Wyman-Changeux model after 50 years.” Annu Rev Biophys 41: 103–133.

Chrisman, I. M., M. D. Nemetchek, I. M. S. de Vera, J. Shang, Z. Heidari, Y. Long, H. Reyes-Caballero, R. Galindo-Murillo, T. E. Cheatham, 3rd, A. L. Blayo, Y. Shin, J. Fuhrmann, P. R. Griffin, T. M. Kamenecka, D. J. Kojetin and T. S. Hughes (2018). “Defining a conformational ensemble that directs activation of PPARgamma.” Nat Commun 9(1): 1794.

Cimmperman, P., L. Baranauskiene, S. Jachimoviciute, J. Jachno, J. Torresan, V. Michailoviene, J. Matuliene, J. Sereikaite, V. Bumelis and D. Matulis (2008). “A quantitative model of thermal stabilization and destabilization of proteins by ligands.” Biophys J 95(7): 3222–3231.

Clark, L. D., I. Dikiy, K. Chapman, K. E. Rodstrom, J. Aramini, M. V. LeVine, G. Khelashvili, S. G. Rasmussen, K. H. Gardner and D. M. Rosenbaum (2017). “Ligand modulation of sidechain dynamics in a wild-type human GPCR.” Elife 6.

Dosa, P. I. and E. A. Amin (2016). “Tactical Approaches to Interconverting GPCR Agonists and Antagonists.” J Med Chem 59(3): 810–840.

Eddy, M. T., M. Y. Lee, Z. G. Gao, K. L. White, T. Didenko, R. Horst, M. Audet, P. Stanczak, K. M. McClary, G. W. Han, K. A. Jacobson, R. C. Stevens and K. Wuthrich (2018). “Allosteric Coupling of Drug Binding and Intracellular Signaling in the A2A Adenosine Receptor.” Cell 172(1-2): 68–80 e12.

Emsley, P. and K. Cowtan (2004). “Coot: model-building tools for molecular graphics.” Acta Crystallogr D Biol Crystallogr 60(Pt 12 Pt 1): 2126–2132.

Fraser, J. S., M. W. Clarkson, S. C. Degnan, R. Erion, D. Kern and T. Alber (2009). “Hidden alternative structures of proline isomerase essential for catalysis.” Nature 462(7273): 669–673.

Fraser, J. S., H. van den Bedem, A. J. Samelson, P. T. Lang, J. M. Holton, N. Echols and T. Alber (2011). “Accessing protein conformational ensembles using room-temperature X-ray crystallography.” Proc Natl Acad Sci U S A 108(39): 16247–16252.

Fujioka, M. and N. Omori (2012). “Subtleties in GPCR drug discovery: a medicinal chemistry perspective.” Drug Discov Today 17(19-20): 1133–1138.

Ge, K., M. Guermah, C. X. Yuan, M. Ito, A. E. Wallberg, B. M. Spiegelman and R. G. Roeder (2002). “Transcription coactivator TRAP220 is required for PPAR gamma 2-stimulated adipogenesis.” Nature 417(6888): 563–567.

Grutsch, S., S. Bruschweiler and M. Tollinger (2016). “NMR Methods to Study Dynamic Allostery.” PLoS Comput Biol 12(3): e1004620.

Hughes, T. S., M. J. Chalmers, S. Novick, D. S. Kuruvilla, M. R. Chang, T. M. Kamenecka, M. Rance, B. A. Johnson, T. P. Burris, P. R. Griffin and D. J. Kojetin (2012). “Ligand and receptor dynamics contribute to the mechanism of graded PPARgamma agonism.” Structure 20(1): 139–150.

Hughes, T. S., P. K. Giri, I. M. de Vera, D. P. Marciano, D. S. Kuruvilla, Y. Shin, A. L. Blayo, T. M. Kamenecka, T. P. Burris, P. R. Griffin and D. J. Kojetin (2014). “An alternate binding site for PPARgamma ligands.” Nat Commun 5: 3571.

Hughes, T. S., J. Shang, R. Brust, I. M. S. de Vera, J. Fuhrmann, C. Ruiz, M. D. Cameron, T. M. Kamenecka and D. J. Kojetin (2016). “Probing the Complex Binding Modes of the PPARgamma Partial Agonist 2-Chloro-N-(3-chloro-4-((5-chlorobenzo[d]thia-zol-2-yl)thio)phenyl)-4-(trifluorome thyl)benzenesulfonamide (T2384) to Orthosteric and Allosteric Sites with NMR Spectroscopy.” J Med Chem 59(22): 10335–10341.

Isogai, S., X. Deupi, C. Opitz, F. M. Heydenreich, C. J. Tsai, F. Brueckner, G. F. Schertler, D. B. Veprintsev and S. Grzesiek (2016). “Backbone NMR reveals allosteric signal transduction networks in the beta1-adrenergic receptor.” Nature 530(7589): 237–241.

Jang, J. Y., M. Koh, H. Bae, D. R. An, H. N. Im, H. S. Kim, J. Y. Yoon, H. J. Yoon, B. W. Han, S. B. Park and S. W. Suh (2017). “Structural basis for differential activities of enantiomeric PPARgamma agonists: Binding of S35 to the alternate site.” Biochim Biophys Acta Proteins Proteom 1865(6): 674–681.

Johnson, B. A. (2004). “Using NMRView to visualize and analyze the NMR spectra of macromolecules.” Methods Mol Biol 278: 313–352.

Johnson, B. A., E. M. Wilson, Y. Li, D. E. Moller, R. G. Smith and G. Zhou (2000). “Ligand-induced stabilization of PPARgamma monitored by NMR spectroscopy: implications for nuclear receptor activation.” J Mol Biol 298(2): 187–194.

Kar, G., O. Keskin, A. Gursoy and R. Nussinov (2010). “Allostery and population shift in drug discovery.” Curr Opin Pharmacol 10(6): 715–722.

Keedy, D. A., L. R. Kenner, M. Warkentin, R. A. Woldeyes, J. B. Hopkins, M. C. Thompson, A. S. Brewster, A. H. Van Ben-schoten, E. L. Baxter, M. Uervirojnangkoorn, S. E. McPhillips, J. Song, R. Alonso-Mori, J. M. Holton, W. I. Weis, A. T. Brunger, S. M. Soltis, H. Lemke, A. Gonzalez, N. K. Sauter, A. E. Cohen, H. van den Bedem, R. E. Thorne and J. S. Fraser (2015). “Mapping the conformational landscape of a dynamic enzyme by multitemperature and XFEL crystallography.” Elife 4.

Kenakin, T. (2004). “Principles: receptor theory in pharmacology.” Trends Pharmacol Sci 25(4): 186–192.

Kenakin, T. (2014). “What is pharmacological ‘affinity’? Relevance to biased agonism and antagonism.” Trends Pharmacol Sci 35(9): 434–441.

Kenakin, T. (2017). “A Scale of Agonism and Allosteric Modulation for Assessment of Selectivity, Bias, and Receptor Mutation.” Mol Pharmacol 92(4): 414–424.

Kim, T. H., K. Y. Chung, A. Manglik, A. L. Hansen, R. O. Dror, T. J. Mildorf, D. E. Shaw, B. K. Kobilka and R. S. Prosser (2013). “The role of ligands on the equilibria between functional states of a G protein-coupled receptor.” J Am Chem Soc 135(25): 9465–9474.

Kofuku, Y., T. Ueda, J. Okude, Y. Shiraishi, K. Kondo, M. Maeda, H. Tsujishita and I. Shimada (2012). “Efficacy of the beta(2)-adrenergic receptor is determined by conformational equilibrium in the transmembrane region.” Nat Commun 3: 1045.

Kofuku, Y., T. Ueda, J. Okude, Y. Shiraishi, K. Kondo, T. Mizumura, S. Suzuki and I. Shimada (2014). “Functional dynamics of deuterated beta2-adrenergic receptor in lipid bilayers revealed by NMR spectroscopy.” Angew Chem Int Ed Engl 53(49): 13376–13379.

Kojetin, D. J. and T. P. Burris (2013). “Small molecule modulation of nuclear receptor conformational dynamics: implications for function and drug discovery.” Mol Pharmacol 83(1): 1–8.

Laghezza, A., L. Piemontese, C. Cerchia, R. Montanari, D. Capelli, M. Giudici, M. Crestani, P. Tortorella, F. Peiretti, G. Pochetti, A. Lavecchia and F. Loiodice (2018). “Identification of the First PPARalpha/gamma Dual Agonist Able To Bind to Canonical and Alternative Sites of PPARgamma and To Inhibit Its Cdk5-Mediated Phosphorylation.” source J Med Chem 61(18): 8282–8298.

Lee, M. A., L. Tan, H. Yang, Y. G. Im and Y. J. Im (2017). “Structures of PPARgamma complexed with lobeglitazone and pioglitazone reveal key determinants for the recognition of antidiabetic drugs.” Sci Rep 7(1): 16837.

Leff, P. (1995). “The two-state model of receptor activation.” Trends Pharmacol Sci 16(3): 89–97.

Liberato, M. V., A. S. Nascimento, S. D. Ayers, J. Z. Lin, A. Cvoro, R. L. Silveira, L. Martinez, P. C. Souza, D. Saidemberg, T. Deng, A. A. Amato, M. Togashi, W. A. Hsueh, K. Phillips, M. S. Palma, F. A. Neves, M. S. Skaf, P. Webb and I. Polikarpov (2012). “Medium chain fatty acids are selective peroxisome proliferator activated receptor (PPAR) gamma activators and pan-PPAR partial agonists.” PLoS One 7(5): e36297.

Liu, J. J., R. Horst, V. Katritch, R. C. Stevens and K. Wuthrich (2012). “Biased signaling pathways in beta2-adrenergic receptor characterized by 19F-NMR.” Science 335(6072): 1106–1110.

Maehle, A. H., C. R. Prull and R. F. Halliwell (2002). “The emergence of the drug receptor theory.” Nat Rev Drug Discov 1(8): 637–641.

Manglik, A., T. H. Kim, M. Masureel, C. Altenbach, Z. Yang, D. Hilger, M. T. Lerch, T. S. Kobilka, F. S. Thian, W. L. Hubbell, R. S. Prosser and B. K. Kobilka (2015). “Structural Insights into the Dynamic Process of beta2-Adrenergic Receptor Signaling.” Cell 161(5): 1101–1111.

Marciano, D. P., D. S. Kuruvilla, S. V. Boregowda, A. Asteian, T. S. Hughes, R. Garcia-Ordonez, C. A. Corzo, T. M. Khan, S. J. Novick, H. Park, D. J. Kojetin, D. G. Phinney, J. B. Bruning, T. M. Kamenecka and P. R. Griffin (2015). “Pharmacological repression of PPARgamma promotes osteogenesis.” Nat Commun 6: 7443.

Mauldin, R. V., M. J. Carroll and A. L. Lee (2009). “Dynamic dysfunction in dihydrofolate reductase results from antifolate drug binding: modulation of dynamics within a structural state.” Structure 17(3): 386–394.

McCoy, A. J., R. W. Grosse-Kunstleve, P. D. Adams, M. D. Winn, L. C. Storoni and R. J. Read (2007). “Phaser crystallographic software.” J Appl Crystallogr 40(Pt 4): 658–674.

Monod, J., J. Wyman and J. P. Changeux (1965). “On the Nature of Allosteric Transitions: A Plausible Model.” J Mol Biol 12: 88–118.

Mosure, S., J. Shang, J. Eberhardt, R. Brust, J. Zheng, P. R. Griffin, S. Forli and D. J. Kojetin (2019). “Structural basis of altered potency and efficacy displayed by a major in vivo metabolite of the anti-diabetic PPARgamma drug pioglitazone.” J Med Chem.

Nolte, R. T., G. B. Wisely, S. Westin, J. E. Cobb, M. H. Lambert, R. Kurokawa, M. G. Rosenfeld, T. M. Willson, C. K. Glass and M. V. Milburn (1998). “Ligand binding and co-activator assembly of the peroxisome proliferator-activated receptor-gamma.” Nature 395(6698): 137–143.

Nygaard, R., Y. Zou, R. O. Dror, T. J. Mildorf, D. H. Arlow, A. Manglik, A. C. Pan, C. W. Liu, J. J. Fung, M. P. Bokoch, F. S. Thian, T. S. Kobilka, D. E. Shaw, L. Mueller, R. S. Prosser and B. K. Kobilka (2013). “The dynamic process of beta(2)-adrenergic receptor activation.” Cell 152(3): 532–542.

Okude, J., T. Ueda, Y. Kofuku, M. Sato, N. Nobuyama, K. Kondo, Y. Shiraishi, T. Mizumura, K. Onishi, M. Natsume, M. Maeda, H. Tsujishita, T. Kuranaga, M. Inoue and I. Shimada (2015). “Identification of a Conformational Equilibrium That Determines the Efficacy and Functional Selectivity of the mu-Opioid Receptor.” Angew Chem Int Ed Engl 54(52): 15771–15776.

Peng, J. W. (2009). “Communication breakdown: protein dynamics and drug design.” Structure 17(3): 319–320.

Prosser, R. S., L. Ye, A. Pandey and A. Orazietti (2017). “Activation processes in ligand-activated G protein-coupled receptors: A case study of the adenosine A2A receptor.” Bioessays 39(9).

Samama, P., S. Cotecchia, T. Costa and R. J. Lefkowitz (1993). “A mutation-induced activated state of the beta 2-adrenergic receptor. Extending the ternary complex model.” J Biol Chem 268(7): 4625–4636.

Savkur, R. S. and T. P. Burris (2004). “The coactivator LXXLL nuclear receptor recognition motif.” J Pept Res 63(3): 207–212.

Shang, J., R. Brust, S. A. Mosure, J. Bass, P. Munoz-Tello, H. Lin, T. S. Hughes, M. Tang, Q. Ge, T. M. Kamenekca and D. J. Kojetin (2018). “Cooperative cobinding of synthetic and natural ligands to the nuclear receptor PPARgamma.” Elife 7.

Solt, A. S., M. J. Bostock, B. Shrestha, P. Kumar, T. Warne, C. G. Tate and D. Nietlispach (2017). “Insight into partial agonism by observing multiple equilibria for ligand-bound and Gs-mimetic nanobody-bound beta1-adrenergic receptor.” Nat Commun 8(1): 1795.

Sounier, R., C. Mas, J. Steyaert, T. Laeremans, A. Manglik, W. Huang, B. K. Kobilka, H. Demene and S. Granier (2015). “Propagation of conformational changes during mu-opioid receptor activation.” Nature 524(7565): 375–378.

Staus, D. P., R. T. Strachan, A. Manglik, B. Pani, A. W. Kahsai, T. H. Kim, L. M. Wingler, S. Ahn, A. Chatterjee, A. Masoudi, A. C. Kruse, E. Pardon, J. Steyaert, W. I. Weis, R. S. Prosser, B. K. Kobilka, T. Costa and R. J. Lefkowitz (2016). “Allosteric nanobodies reveal the dynamic range and diverse mechanisms of G-protein-coupled receptor activation.” Nature 535(7612): 448–452.

Susac, L., M. T. Eddy, T. Didenko, R. C. Stevens and K. Wuthrich (2018). “A2A adenosine receptor functional states characterized by (19)F-NMR.” Proc Natl Acad Sci U S A 115(50): 12733–12738.

Tyka, M. D., D. A. Keedy, I. Andre, F. Dimaio, Y. Song, D. C. Richardson, J. S. Richardson and D. Baker (2011). “Alternate states of proteins revealed by detailed energy landscape mapping.” J Mol Biol 405(2): 607–618.

Weikum, E. R., X. Liu and E. A. Ortlund (2018). “The nuclear receptor superfamily: A structural perspective.” Protein Sci 27(11): 1876–1892.

Weiss, J. M., P. H. Morgan, M. W. Lutz and T. P. Kenakin (1996). “The Cubic Ternary Complex Receptor–Occupancy Model I. Model Description.” J Theor Biol 181(4): 151–167.

Williamson, M. P. (2013). “Using chemical shift perturbation to characterise ligand binding.” Prog Nucl Magn Reson Spectrosc 73: 1–16.

Willson, T. M., J. E. Cobb, D. J. Cowan, R. W. Wiethe, I. D. Correa, S. R. Prakash, K. D. Beck, L. B. Moore, S. A. Kliewer and J. M. Lehmann (1996). “The structure-activity relationship between peroxisome proliferator-activated receptor gamma agonism and the antihyperglycemic activity of thiazolidinediones.” J Med Chem 39(3): 665–668.

Winn, M. D., C. C. Ballard, K. D. Cowtan, E. J. Dodson, P. Emsley, P. R. Evans, R. M. Keegan, E. B. Krissinel, A. G. Leslie, A. McCoy, S. J. McNicholas, G. N. Murshudov, N. S. Pannu, E. A. Potterton, H. R. Powell, R. J. Read, A. Vagin and K. S. Wilson (2011). “Overview of the CCP4 suite and current developments.” Acta Crystallogr D Biol Crystallogr 67(Pt 4): 235–242.

Ye, L., N. Van Eps, M. Zimmer, O. P. Ernst and R. S. Prosser (2016). “Activation of the A2A adenosine G-protein-coupled receptor by conformational selection.” Nature 533(7602): 265–268.

