## supplementary data for "Quantitative Structural Assessment of Graded Receptor Agonism"

This file contains:

1 supplementary table
6 supplementary figures

**Supplementary Table S1. X-ray crystallography data collection and refinement statistics.**

| | PPARγ LBD + Darglitazone | PPARγ LBD + CAY10506 | PPARγ LBD + Troglitazone | PPARγ LBD + Ciglitazone | PPARγ LBD + Mitoglitazone | PPARγ LBD + CAY10638 |
|---|---|---|---|---|---|---|
| **Data collection** | | | | | | |
| Space group | C 1 2 1 | C 1 2 1 | C 1 2 1 | C 1 2 1 | C 1 2 1 | C 1 2 1 |
| Cell dimensions | | | | | | |
| *a*, *b*, *c* (Å) | 93.36, 61.74, 119.47 | 92.20, 59.49, 116.73 | 92.81, 62.07, 118.76 | 92.39, 59.96, 117.61 | 92.93, 60.35, 118.11 | 92.86, 60.60, 117.66 |
| *α*, *β*, *γ* (°) | 90.00, 102.72, 90.00 | 90.00, 103.55, 90.00 | 90.00 102.10 90.00 | 90.00, 103.02, 90.00 | 90.00, 103.14, 90.00 | 90.00, 103.14, 90.00 |
| Resolution | 41.01-1.95 (2.02-1.95) | 45.48-2.45 (2.54-2.45) | 44.95-3.10 (3.21-3.10) | 49.90-2.78 (2.88-2.78) | 48.30-2.52 (2.61-2.52) | 50.34-2.15 (2.23-2.15) |
| $R_{pim}$ | 0.037 (0.525) | 0.040 (0.280) | 0.0726 (0.378) | 0.059 (0.457) | 0.036 (0.303 | 0.055 (0.409) |
| I / σ(I) | 8.62 (1.26) | 12.38 (2.89) | 8.48 (2.20) | 8.13 (1.71) | 11.79 (2.54) | 5.84 (1.34) |
| CC1/2 in highest shell | 0.611 | 0.811 | 0.717 | 0.642 | 0.814 | 0.805 |
| Completeness (%) | 96.06 (94.82) | 97.30 (96.95) | 98.55 (97.71) | 99.77 (99.94) | 99.92 (99.77) | 96.42 (92.89) |
| Redundancy | 1.7 (1.6) | 2.0 (2.0) | 1.8 (1.8) | 1.9 (1.9) | 2.0 (2.0) | 1.7 (1.7) |
| **Refinement** | | | | | | |
| Resolution (Å) | 1.95 | 2.45 | 3.10 | 2.78 | 2.52 | 2.15 |
| No. of unique reflections | 46650 | 22240 | 12003 | 15994 | 21743 | 33678 |
| $R_{work}/R_{free}$ (%) | 22.6/27.0 | 19.6/24.7 | 19.4/27.7 | 24.0/30.1 | 19.2/24.5 | 21.9/26.2 |
| No. of atoms | | | | | | |
| Protein | 4147 | 4080 | 4141 | 4169 | 4098 | 4132 |
| Water | 423 | 185 | 0 | 15 | 158 | 226 |
| *B*-factors | | | | | | |
| Protein | 27.90 | 31.05 | 25.38 | 65.65 | 31.90 | 30.45 |
| Ligand | 30.67 | 39.11 | 30.00 | 87.49 | 41.61 | 39.01 |
| Water | 32.29 | 30.12 | n/a | 56.79 | 30.57 | 31.88 |
| Root mean square deviations | | | | | | |
| Bond lengths (Å) | 0.008 | 0.009 | 0.011 | 0.010 | 0.008 | 0.009 |
| Bond angles (°) | 0.92 | 1.03 | 1.16 | 1.27 | 1.02 | 1.09 |
| Ramachandran favored (%) | 95.66 | 95.98 | 89.80 | 92.16 | 97.39 | 95.41 |
| Ramachandran outliers (%) | 1.18 | 0.80 | 1.93 | 0.93 | 0.40 | 1.30 |
| PDB accession code | 6DGL | 6DGQ | 6DGO | 6O68 | 6O67 | 6DGR |

**Values in parentheses indicate highest resolution shell.*

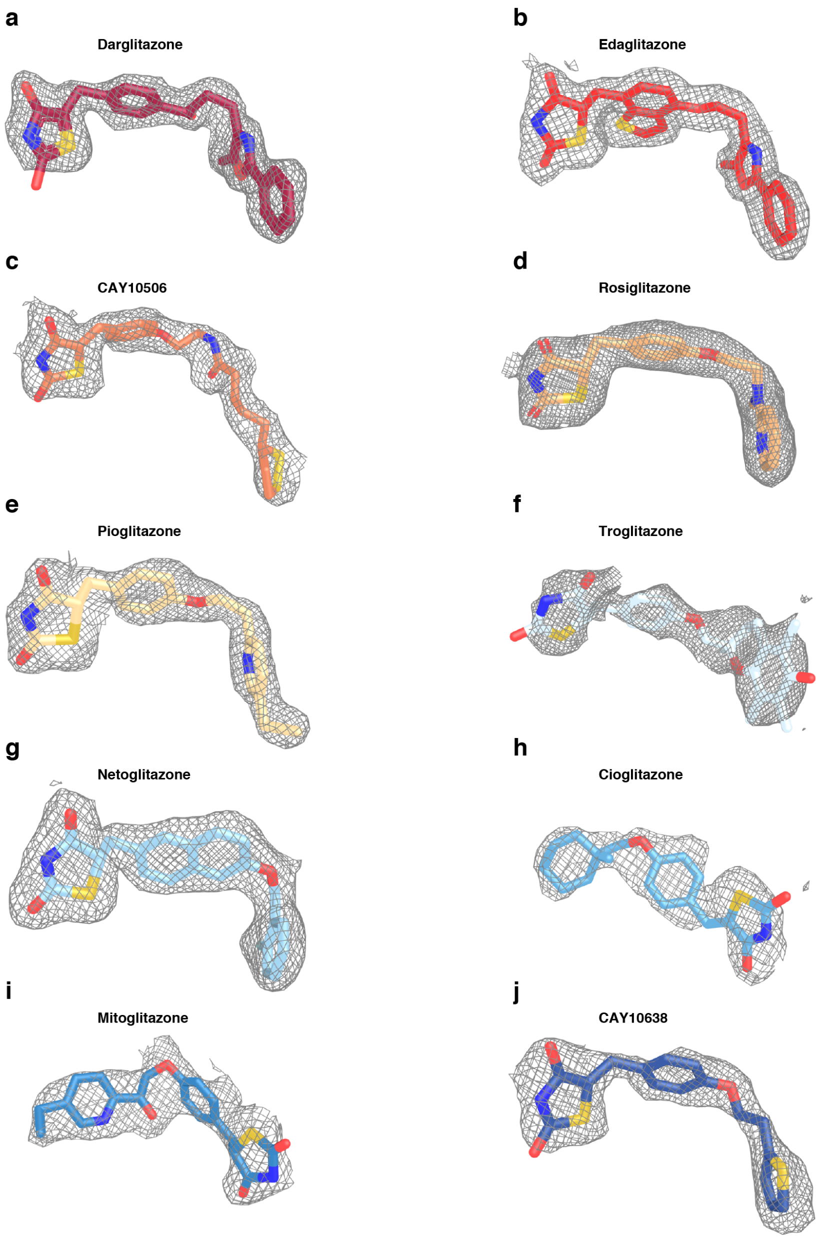

**Supplementary Figure S1.** TZD ligand density in the crystal structures. Omit maps ($2F_O–F_C$, contoured at 1σ) for PPARγ LBD bound to (**a**) Darglitazone (PDB 6DGL), (**b**) Edaglitazone (PDB 5UGM), (**c**) CAY10506 (PDB 6DGQ), (**d**) Rosiglitazone (PDB 4EMA), (**e**) Pioglitazone (PDB 5Y2O), (**f**) Troglitazone (PDB 6DGO), (**g**) Netoglitazone (PDB 3B0Q), (**h**) Cioglitazone (PDB 6O68), (**i**) Mitoglitazone (PDB 6O67), and (**j**) CAY10638 (PDB 6DGR).

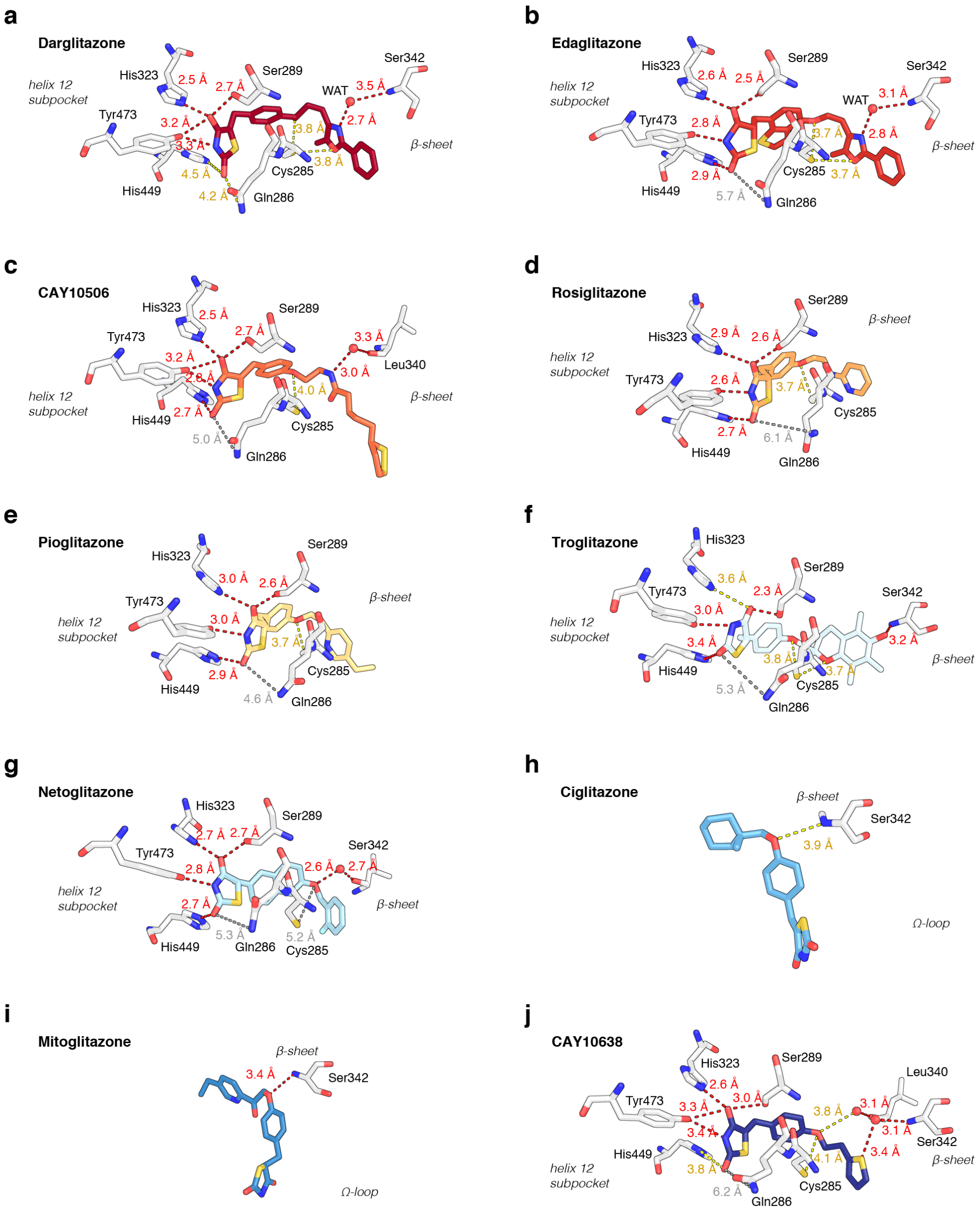

**Supplementary Figure S2.** Ligand hydrogen bond and electrostatic contacts in the crystal structures. Distances are represented by dotted lines colored colored by interaction strength: strong, <3.5Å (red); moderate, 3.5–4.5Å (yellow); weak, >4.5Å (gray) for PPARγ LBD bound to (**a**) Darglitazone (PDB 6DGL), (**b**) Edaglitazone (PDB 5UGM), (**c**) CAY10506 (PDB 6DGQ), (**d**) Rosiglitazone (PDB 4EMA), (**e**) Pioglitazone (PDB 5Y2O), (**f**) Troglitazone (PDB 6DGO), (**g**) Netoglitazone (PDB 3B0Q), (**h**) Cioglitazone (PDB 6O68), (**i**) Mitoglitazone (PDB 6O67), and (**j**) CAY10638 (PDB 6DGR).

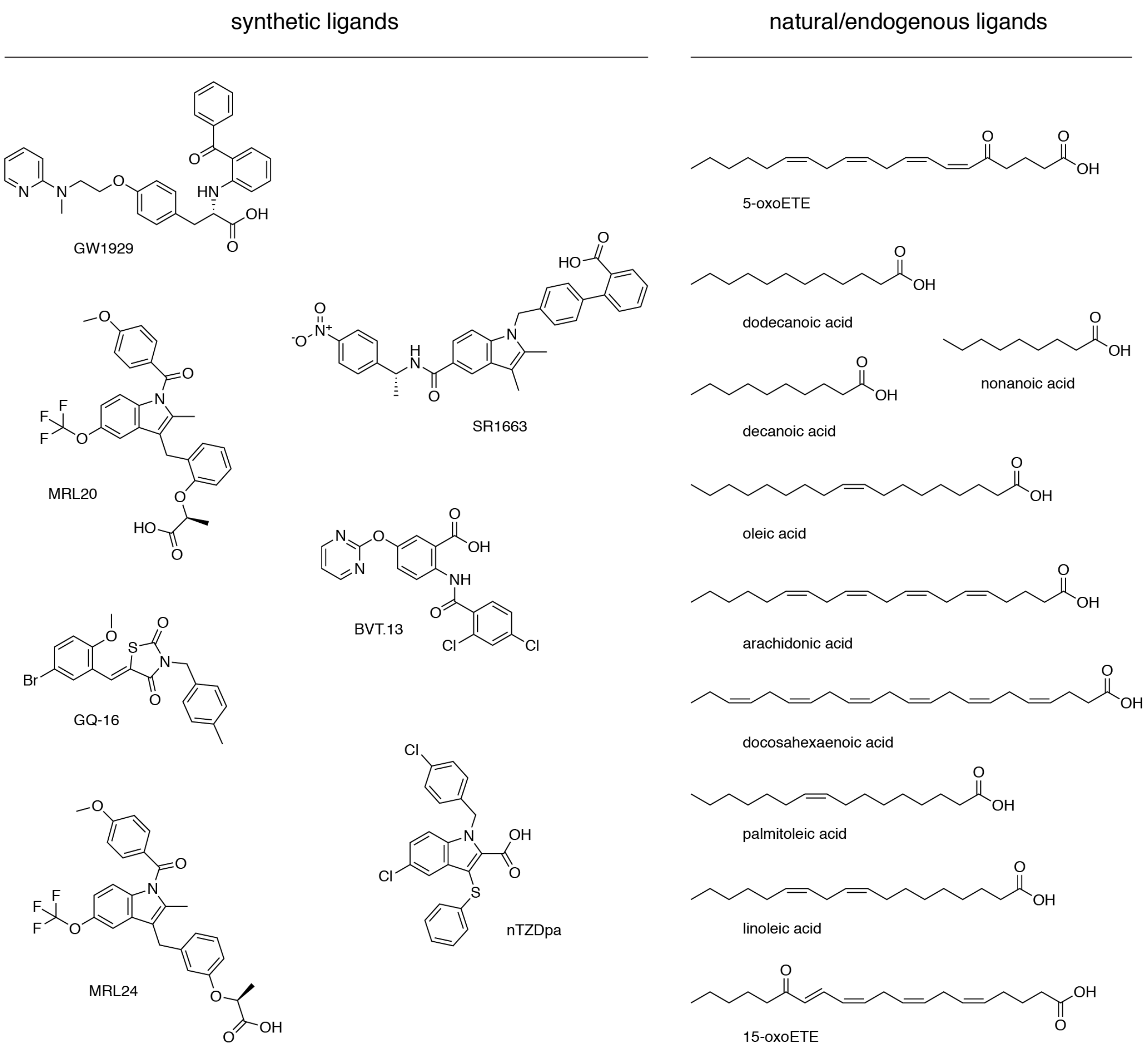

**Supplementary Figure S3.** Chemical structures of the diverse ligand set.

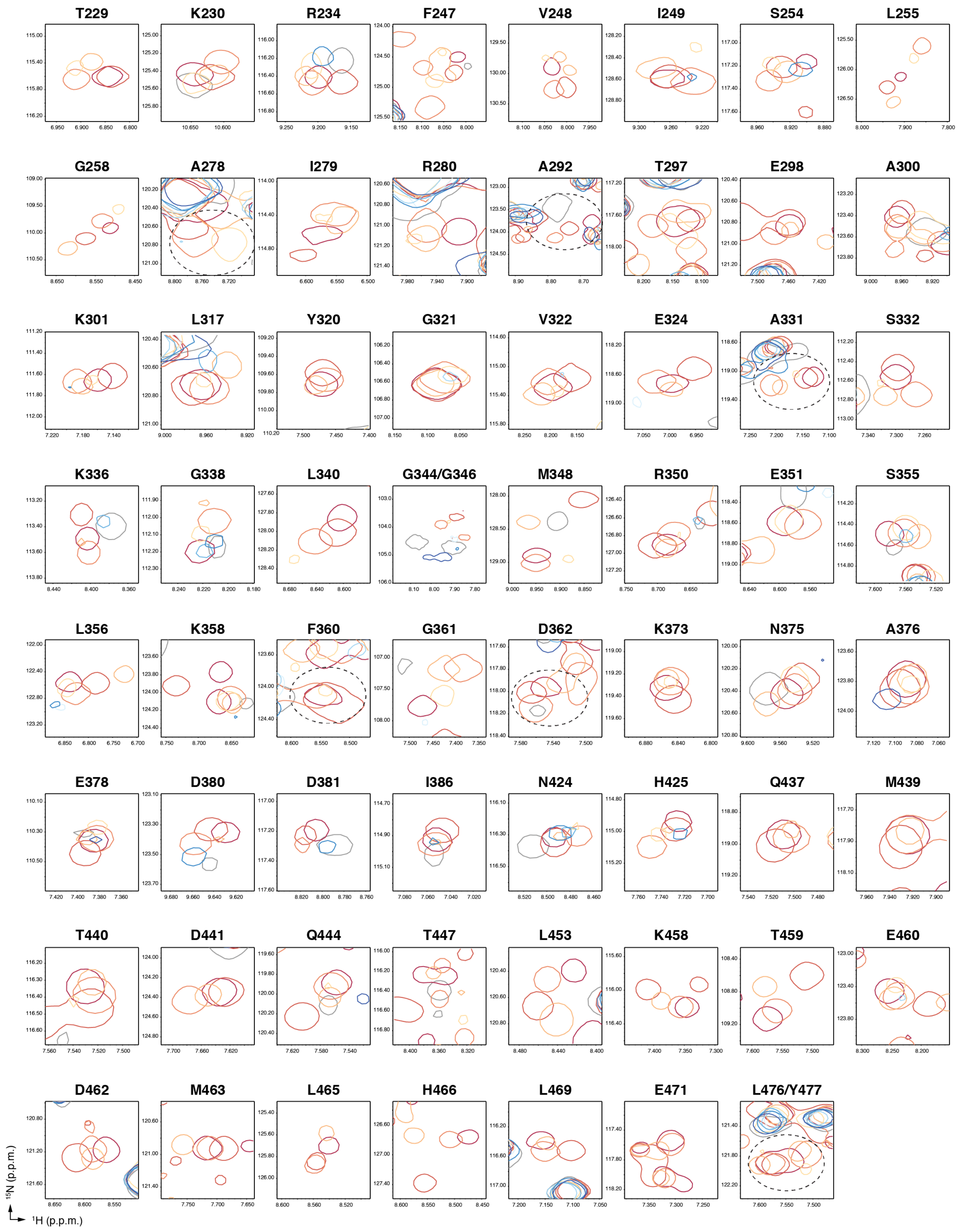

**Supplementary Figure S4.** Other residues that show NMR peak line broadening for less potent ligands in the TZD series.

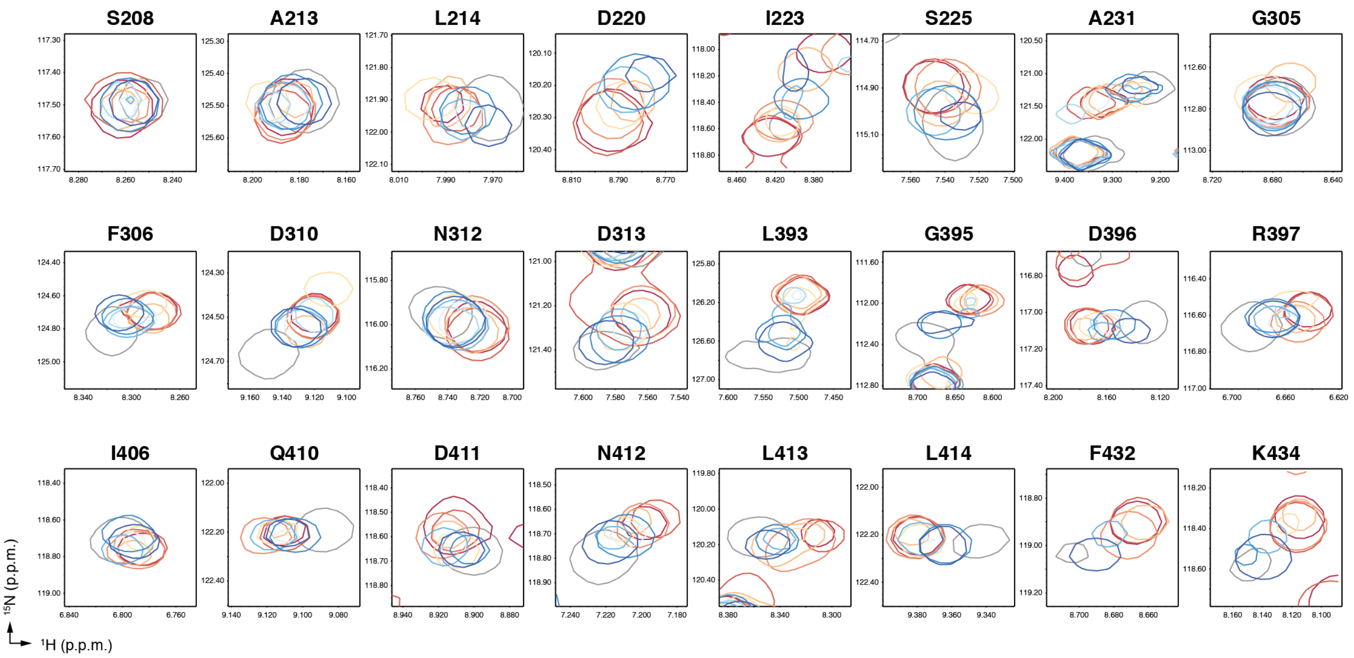

**Supplementary Figure S5.** Other residues that show co-linear NMR peak shifting in the TZD series.

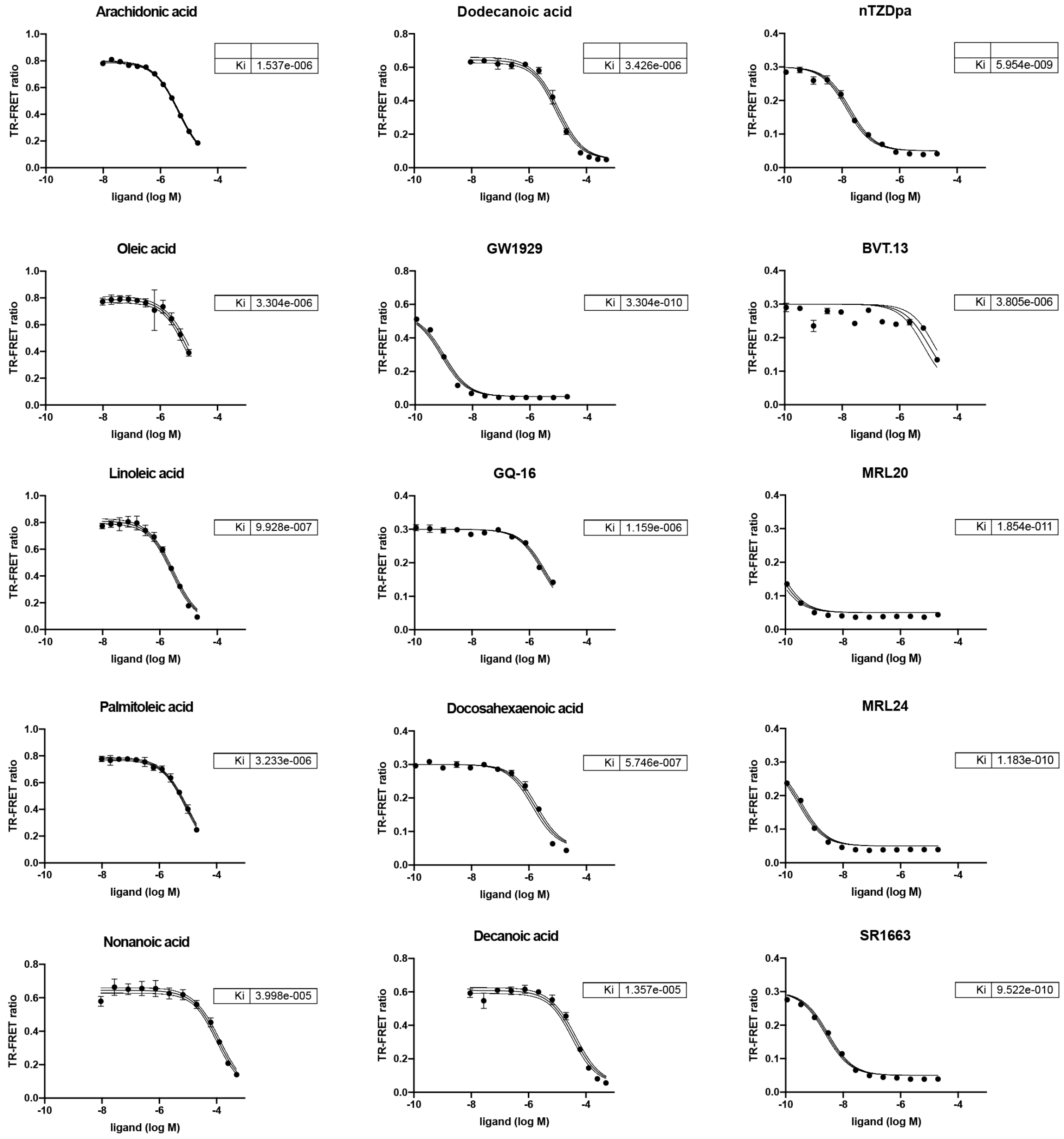

**Supplementary Figure S6.** TR-FRET assay to determine ligand affinity ($K_i$ values) using a fluorescent tracer ligand fit to the Cheng-Prusoff inhibitor constant equation; error bars, mean ± s.d. (n=3); fitted line (solid) shown along with the 95% confidence interval (dotted line).
